# Metabolic alterations drive inflammatory phenotypes in CHIP-associated heart failure

**DOI:** 10.1101/2024.07.30.605744

**Authors:** Siavash Mansouri, Inga Hemmerling, Golnaz Hesami, Sebastian Cremer, Klara Kirschbaum, Stephan Klatt, Fatemeh Behjati, Marcel H. Schulz, Xue Li, Marina Scheller, Carsten Müller-Tidow, Philipp Friedrich Arndt, Stefan Günther, Manju Padmasekar, Mario Looso, Michael A. Rieger, Stefan Kuhnert, Claus F. Vogelmeier, Robert Bals, Ingrid Fleming, Andreas Zeiher, Werner Seeger, Florian Leuschner, Rajkumar Savai, Stefanie Dimmeler, Soni S Pullamsetti

## Abstract

Mutations in DNA methyltransferase 3 alpha *(DNMT3A)* are the most frequent driver of clonal hematopoiesis of indeterminate potential (CHIP), and associated with higher risk of cardiovascular disease and pro-inflammatory activation of immune cells. Here, we investigated the mechanisms underlying DNMT3A CHIP-associated inflammatory phenotypes in macrophages. We show that monocytes of *DNMT3A* CHIP-driver mutation carriers are associated with DNA hypomethylation of succinate dehydrogenase A (SDHA) and an altered tricarboxylic acid cycle metabolite profile. Silencing of DNMT3A in monocytes increased SDHA and elevated mitochondria complex II activity. The secreted complex II product, malate, further increased inflammatory activation in wild type monocytes to further augment inflammation in a paracrine manner. Pharmacological inhibition of SDHA (using dimethyl malonate) in mice harboring *DNMT3A* mutations in hematopoietic stem cells ameliorated the inflammatory response and improved cardiac function after myocardial infarction. Thus, interfering with the altered metabolic state may provide a new therapeutic option to dampen inflammatory activation in DNMT3A CHIP carrying patients.

## Introduction

Clonal hematopoiesis of indeterminate potential (CHIP) is induced by single recurrent somatic mutations in hematopoietic stem cells (HSC) in the bone marrow, which lead to clonal dominance. CHIP occurs frequently in an age-dependent fashion and has emerged as a major risk factor associated with several cardiovascular diseases (CVD)^1^. Sequencing studies in large cohorts revealed that CHIP driver mutations in the genes *DNMT3A*, *TET2*, *ASXL1*, and *JAK2* are associated with an increased risk of heart and lung disease, stroke and increased mortality^2,3,4^. Importantly, the prevalence of CHIP is 4-fold higher in individuals with myocardial infarction^3^, and the most commonly mutated CHIP driver genes i.e. *TET2* and *DNMT3A*, are associated with the progression and poor prognosis of chronic heart failure ^5,6^. A dose-response correlation of the size of the mutated blood cell clone with patient survival suggested a functional role of mutated blood cells and the progression of heart failure ^6,7^. The causal involvement of CHIP-mutated blood cells was confirmed in murine models, showing that inactivation of *Tet2* in hematopoietic cells exacerbated the development of cardiovascular diseases, such as atherosclerosis and heart failure, and also impaired lung function in models of cigarette smoke exposure ^4,8,9^. Gene editing of DNMT3A in hematopoietic stem cells additionally impaired heart function after angiotensin II infusion ^10^. Because of the high prevalence and the negative impact of CHIP on human health, there is an unmet need for effective treatment options.

Recent studies have suggested that the detrimental effects of CHIP mutations are mediated by a pro-inflammatory activation of immune cells. Specifically, *TET2 or DNMT3A* mutations increase pro-inflammatory cytokine release by myeloid cells. For example, individuals carrying *TET2* mutations present higher levels of circulating IL-8 and larger numbers of non-classical inflammatory monocytes, likely contributing to the development of CVD^3^. In line with this, atherosclerotic *Tet-2* deficient mice have an increased plaque size and higher numbers of inflammatory leukocytes, in particular, IL-1β producing macrophages which act as potential amplifiers of disease pathology^11^. Furthermore, single cell RNA sequencing of peripheral blood-derived monocytes from individuals with heart failure or aortic valve stenosis who carry TET2 or *DNMT3A* mutations exhibit increased expression of pro-inflammatory cytokines including IL-6, IL-1β ^12,13^. CHIP carrying patient monocytes also presented an augmented gene signature known to promote adherence to endothelial cells as well as interactions with T cells^13,14^. The specific molecular mechanism(s) that drives pro-inflammatory gene expression in response to mutations of the DNA methylation regulatory genes TET2 or DNMT3A, however, is unclear.

The growing field of immunometabolism emphasizes the importance of cellular metabolism for immune cell health and function ^15-18^. Recent studies have revealed that innate immune cells such as macrophages possess distinct metabolic characteristics that correlate with their functional state. The molecular processes that translate such metabolic reprogramming into altered immune-associated gene expression and effector activities are gradually coming to light. For instance, glycolysis contributes to pro-inflammatory M1 macrophage polarization and its inhibition reduces many facets of the inflammatory phenotype, such as phagocytosis, reactive oxygen species (ROS) production, and the secretion of pro-inflammatory cytokines ^19^. Although metabolism has the fundamental regulatory roles in the immune cell activation and inflammatory response ^20^, cellular metabolism has not been studied in CHIP-associated disease.

Here, we report that blood cells and *ex vivo*-cultured macrophages derived from heart failure patients carrying *DNMT3A* CHIP mutations show extensive alterations of cellular metabolism. In detail, DNMT3A CHIP mutations induce the hypomethylation of succinate dehydrogenase (SDHA) leading to the de-repression of SDHA and consequent activation of mitochondrial activity and accumulation of the TCA cycle metabolite malate. Mechanistically, SDHA and malate have a decisive role in the induction of pro-inflammatory phenotype in DNMT3A deficient or mutant macrophages. Furthermore, interfering with SDHA activity using dimethyl malonate reversed DNMT3A mutation-induced inflammatory activation and improved cardiac function in mice that conditionally expresses human *DNMT3A* cDNA carrying the R882H mutation, one of the major mutational hotspot in clonal hematopoiesis ^21^. Thus, being at the crossroads of epigenetics and immunity, metabolism and metabolic modulation can provide a unique therapeutic opportunity for CHIP-associated diseases.

## Results

### DNMT3A CHIP carriers show SDHA hypomethylation and accumulation of TCA metabolites

Given the functional role of DNMT3A in *de novo* DNA methylation in patients with COPD (Chronic obstructive pulmonary disease) and cardiovascular disease ^8,22,23^, we examined whether there are hypomethylated regions common to both diseases (Figure 1A and Table S1). Therefore, we assessed DNA methylation in peripheral blood cells from heart failure and COPD patients with and without *DNMT3A* CHIP mutations (Table S2) using the Infinium Human-Methylation EPIC BeadChip (850k) (WG-317, Illumina) which covers more than 850.000 CpG sites throughout the whole genome. Interestingly, when promoter methylation levels were ranked on the basis of β-value, fold change and differential methylation p-value, promoter DNA methylation of none of the inflammatory genes has been altered (Table S3A-S3B), but the DNA methylation of the promoter of succinate dehydrogenase complex flavoprotein subunit A (*SDHA)* was found lower in DNMT3A CHIP patients, in both disease indications (Figures 1B and 1C; Table S4). SDHA catalyzes the oxidation of succinate to fumarate and transfers the resulting electrons to SDHB via the electron transporting coenzyme, flavin adenine dinucleotide (FAD) and serves as a main metabolic regulatory element in the TCA cycle and electron transport chain (ETC) ^24^. To gain insight into potential consequences of SDHA DNA methylation, we performed a metabolome analysis of TCA cycle metabolites in the serum of heart failure and COPD patients with and without *DNMT3A* mutations. As CHIP is highly correlated with aging ^2^, we also measured TCA cycle metabolites in the serum of age-matched healthy donors. Importantly, several of the TCA cycle metabolites especially isocitrate, 2-hydroxyglutarate (2HG), fumarate and malate, were significantly increased in DNMT3A CHIP patients compared to non-CHIP patients and healthy controls (Figure 1D). These data indicated that the *DNMT3A* CHIP mutations can result in SDHA hypomethylation to impact on the TCA cycle. To determine whether the increase in SDHA could account for elevated malate levels, we evaluated the malate/succinate and malate/glutamate ratios, as glutaminolysis is the other possible source of malate ^25^. The analysis revealed a significant increase in the malate/succinate ratio in DNMT3A CHIP patients but not in the malate/glutamate ratio (Figure 1E; Figure S3C). Thus, succinate to malate conversion via SDHA could be responsible for increased malate in DNMT3A CHIP carriers with heart failure or COPD. Together, the results demonstrate a strong association between *DNMT3A* CHIP mutations and altered TCA cycle activity especially *SDHA* hypomethylation and alterations in TCA cycle intermediates.

**Figure 1.**
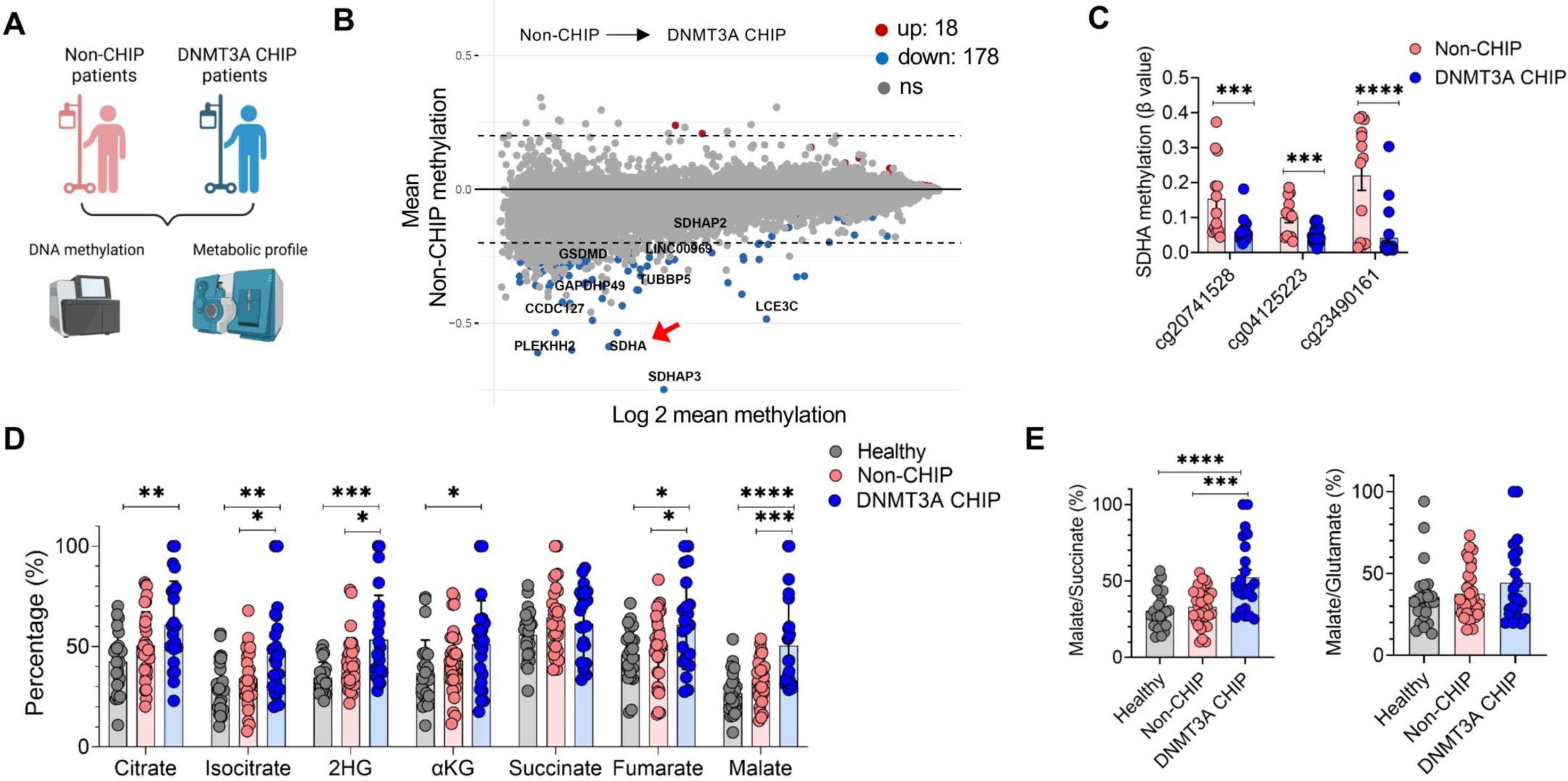
DNA methylation and metabolic profile of CHIP patients with *DNMT3A* mutations. (**A**) Schematic representation of the design of DNA methylation and TCA metabolomics analysis using blood and serum, respectively, from Non-CHIP and DNMT3A CHIP patients. (**B**) MA-plot for mean non-CHIP DNA methylation level against log2 (mean.methylation.DNMT3A/mean.methylation.Non-CHIP) between DNMT3A CHIP (heart failure (HF), n=5; COPD, n=16) and non-CHIP (HF, n=3; COPD, n=10) patients. Top differentially hypomethylated genes according to mean CpG methylation in their promoter region with p<0.01 and fold change ≥0.2 are shown in blue. (**C**) Methylation level of three CG sites cg20741528, cg04125223 and cg23490161 which are located in the *SDHA* promoter between patients with DNMT3A CHIP (HF, n=5; COPD, n=16) and non-CHIP (HF, n=3; COPD, n=10) patients. (**D**) The percentage of serum abundance of TCA metabolites in DNMT3A CHIP (HF, n=8; COPD, n=15) compared with non-CHIP (HF, n=10; COPD, n=22) patients and healthy donors (n=24). (**E**) Ratio of the percentage of malate over succinate and glutamate in patients with *DNMT3A* CHIP mutation (HF, n=8; COPD, n=15) in compared with non-CHIP associated patients (HF, n=10; COPD, n=22) and healthy donors (n=24). Statistical significance was assessed by two-tailed unpaired Student’s *t*-test and one-way ANOVA Tukey’s multiple comparison test. *p <0.05, **p <0.01, ***p< 0.001, ****p< 0.0001.

### DNMT3A CHIP driver mutations augment oxygen consumption rate in macrophages

To gain further insight into the metabolic alterations induced by DNMT3A CHIP driver mutations, we cultured peripheral blood mononuclear cell (PBMC)-derived macrophages from heart failure patients with and without *DNMT3A* CHIP mutations *ex vivo*, and assessed transcriptional changes in metabolic pathways as well as mitochondrial activity. Pathway analysis from RNA-seq data revealed an enrichment of genes associated with oxidative phosphorylation in macrophages of heart failure patients with *DNMT3A* CHIP mutations (Figures 2A and 2B). Consistent with this, we detected a higher oxygen consumption rate (OCR), a functional indicator of oxidative phosphorylation in mitochondria, in macrophages carrying *DNMT3A* CHIP mutations compared to non-CHIP macrophages. Notably, an increase in basal OCR levels and spare capacity was observed, with no significant changes in glycolysis as assessed by the extracellular acidification rate (ECAR) (Figures 2C and 2D). To identify the complexes driving the higher OCR, we analyzed the function of respiratory complexes I–IV by assessing their contribution to oxygen consumption in permeabilized macrophages^26^. This approach identified a clear impact of DNMT3A CHIP mutations on the function of complex II (Figure 2E). There were no consistent differences in mitochondrial ROS production between macrophages with *DNMT3A* CHIP mutations and non-CHIP macrophages (Figure S1A). However, in line with the *SDHA* hypomethylation detected in CHIP DNMT3A patients, *SDHA* mRNA and protein levels were clearly upregulated in *DNMT3A* CHIP mutation carrying macrophages from heart failure patients (Figures 2F and 2G). Next, we confirmed previous studies showing the increased inflammatory phenotype of macrophages, by documenting higher IL1β and IL6 mRNA and protein levels in *DNMT3A* CHIP carriers versus non carriers (Figures 2H-2K). These data indicate that *DNMT3A* CHIP mutations in macrophages cause DNA hypomethylation and upregulation of SDHA, leading to higher complex II activity, that is associated with a pro-inflammatory phenotype to the macrophages.

**Figure 2.**
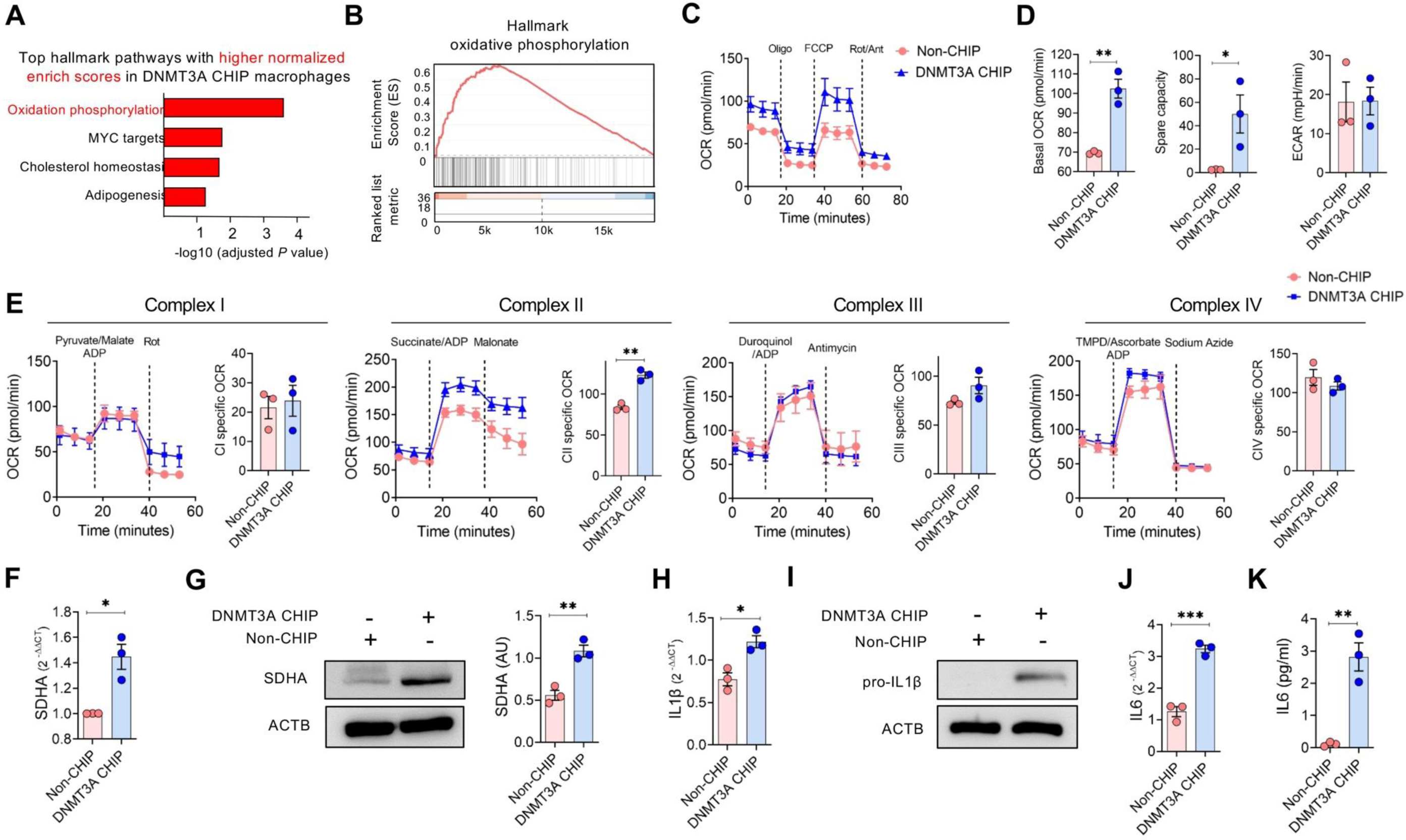
Human DNMT3A CHIP macrophages display a distinct mitochondrial metabolism with the pro-inflammatory phenotype. (**A**) RNA sequencing was performed on PBMC-derived macrophages isolated from HF patients with *DNMT3A* CHIP (n=2) and non-CHIP (n=2) mutation. Top listed pathways with higher normalized enriched scores from the differentially expressed genes from RNA sequencing. (**B**) Gene set enrichment analysis of oxidative phosphorylation genes in PBMC-derived macrophages of HF patients with *DNMT3A* CHIP (n=2) and non-CHIP (n=2) mutations. (**C**) Oxygen consumption rate (OCR) measurement of PBMC-derived macrophages of HF patients with *DNMT3A* CHIP (n=3) and non-CHIP (n=3) mutations. Data are shown as OCR graphs (left) and (**D**) followed by bar graphs of basal respiration, spare capacity and extracellular acidification rate (ECAR) (right). (**E**) OCR measurements of permeabilized PBMC-derived macrophages of HF patients with DNMT3A CHIP (n=3) and non-CHIP Macrophages (n=3) mutations before and after the addition of substrate and associated-complex specific inhibitors, as indicated (dashed lines). Data are shown as OCR graphs (left) and bar graphs of specific complex OCRs (right). (**F**) mRNA expression of *SDHA* in PBMC-derived macrophages of HF patients with *DNMT3A* CHIP (n=3) and non-CHIP (n=3) mutations. (**G**) SDHA protein level followed by the densitometric quantification of relative SDHA expression in PBMC-derived macrophages of HF patients with *DNMT3A* CHIP (n=3) and non-CHIP (n=3) mutations. (**H**) mRNA expression of *IL1β* (n=3) followed by (**I**) protein level of pro-IL1β (n=2) in PBMC-derived macrophages of HF patients with *DNMT3A* CHIP and non-CHIP mutations. (**J, K**) mRNA expression of *IL6* and secretary levels of IL6 in PBMC-derived macrophages of HF patients with *DNMT3A* CHIP (n=3) and non-CHIP (n=3) mutations (*n* = 3). Statistical significance was assessed by a two-tailed unpaired Student’s *t*-test. *p <0.05, **p <0.01, ***p< 0.001

### Loss of DNMT3A in macrophages augments metabolic activity

To assess whether DNMT3A deficiency could mimic *DNMT3A* CHIP mutations to alter TCA cycle activity, we silenced *DNMT3A* in healthy donor-derived macrophages with stealth RNAi against *DNMT3A* (siDNMT3A) and a scramble negative control (siControl) (Figure 3A;Figure S2A). DNMT3A-deficient macrophages expressed higher levels of IL1β and IL6 mRNA and protein (Figures 3B-3E), and demonstrated an enhanced OCR with increased spare capacity, as an indicator of mitochondrial activity (Figure 3F). However, ECAR was similar between DNMT3A-expressing and DNMT3A- deficient macrophages (Figure 3F). Similar to the macrophages carrying *DNMT3A* CHIP mutations, DNMT3A-deficient macrophages demonstrated enhanced complex II activity and an upregulation of *SDHA* (Figures 3G and 3H; Figure S2B). Further, metabolomic analyses also revealed an accumulation of malate and succinate in the supernatant of DNMT3A-deficient macrophages (Figure 3I). DNMT3A- expressing and -deficient macrophages showed the same level of mitochondria ROS production (Figure S2C); the same pattern was also observed between macrophages with *DNMT3A* CHIP mutations and non-CHIP. As CHIP status is mostly associated with chronic inflammation, we challenged DNMT3A deficient macrophages with LPS for 48 hours and then evaluated the mitochondrial metabolism. DNMT3A-deficient macrophages revealed higher OCRs, spare capacity and ECAR level upon LPS stimulation when compared with LPS-stimulated siControl macrophages (Figure S2D). Taken together, these results show that inactivation of *DNMT3A* mimicked the changes to macrophage metabolism induced by *DNMT3A* CHIP mutation to increase the activity of complex II, SDHA expression and pro-inflammatory gene expression.

**Figure 3.**
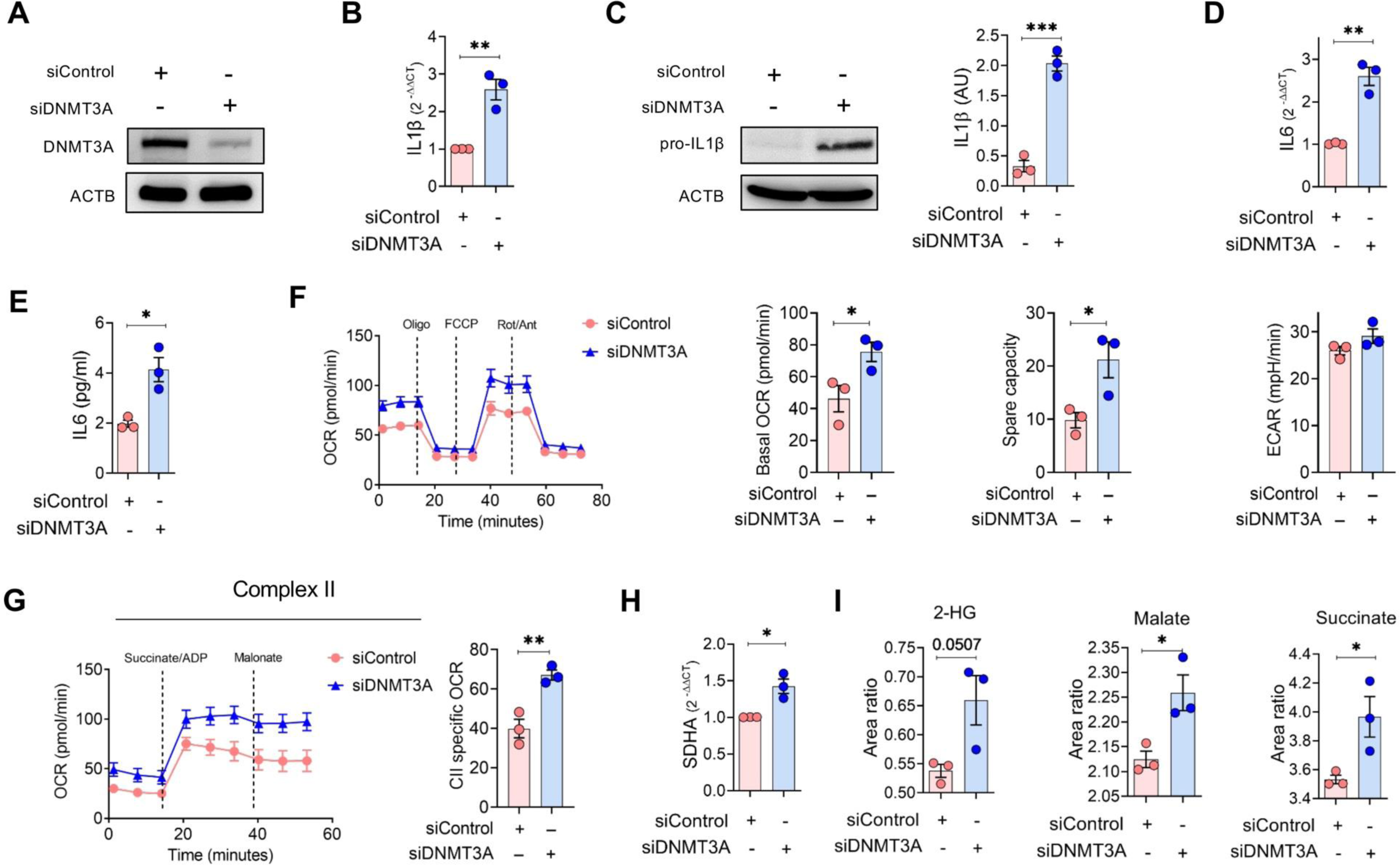
Loss of DNMT3A in macrophages renders a pro-inflammatory and distinct mitochondria metabolism phenotype. (**A**) Protein level of DNMT3A in human PBMC-derived macrophages treated with stealth RNAi against *DNMT3A* (siDNMT3A) and a scramble negative control (siControl) (*n* = 3). (**B**) mRNA expression of *IL1β* and (**C**) protein level of pro-IL1β in DNMT3A deficient PBMC-derived macrophages followed by the densitometric quantification of relative IL1β expression (*n* = 3). (**D**) mRNA expression of *IL6* and (**E**) secretory level of IL6 protein of DNMT3A deficient PBMC-derived macrophages (*n* = 3). (**F**) OCR measurement of DNMT3A deficient human PBMC-derived macrophages (*n* = 3). Data are shown as OCR graphs (left) representative of three independent experiments and summary bar graphs of spare capacity also extracellular acidity rate (ECAR) (right). (**G**) OCR measurements of permeabilized DNMT3A deficient human PBMC-derived macrophages (*n* = 3) before and after the addition of substrate and associated-complex specific inhibitors, as indicated (dashed lines). Data are shown as OCR graphs (left) representative of three independent experiments, and summary bar graphs of complex II specific OCRs (right). (**H**) mRNA expression of *SDHA* in DNMT3A deficient PBMC-derived macrophages (*n* = 3). (**I**) Abundance of 2HG, malate and succinate metabolites in supernatant of DNMT3A deficient human PBMC-derived macrophages (*n* = 3). Statistical significance was assessed by a two-tailed unpaired Student’s *t*-test and. *p <0.05, **p <0.01, ***p< 0.001.

### SDHA/Malate axis regulates pro-inflammatory activation induced by DNMT3A deficiency

To determine whether the pro-inflammatory response observed in DNMT3A CHIP macrophages is a consequence of altered TCA cycle activity accounted for by the upregulation of SDHA and accumulation of malate, we used a siRNA approach to silence *SDHA* in macrophages from heart failure patients carrying *DNMT3A* CHIP mutations (Figure 4A). Notably, the *SDHA* siRNA approach decreased both IL1β and IL6 in macrophages from heart failure patients carrying *DNMT3A* CHIP mutations (Figures 4B and 4C). Similarly, the knock down of both *SDHA* and *DNMT3A* in donor-derived macrophages also reduced IL1β and IL6 levels (Figure 4D; E Figure S3A). The lack of SDHA negatively impacted on the OCR, decreasing of basal respiration and the spare capacity without affecting ECAR in DNMT3A deficient macrophages (Figure 4F). To explore the mechanisms underlying SDHA-induced pro-inflammatory activation, we focused on malate, a downstream product of SDHA activity. Indeed, malate accumulates in macrophages under inflammatory conditions^27^ and in β-glucan-induced trained macrophages^28^. Interestingly, treatment of SDHA/DNMT3A deficient macrophages with malate for 24 hours increased IL1β and IL6 levels in SDHA/DNMT3A-deficient macrophages (Figures 4G and 4H, Figure S3B). These data indicated that SDHA-mediated malate production contributes to the pro-inflammatory response induced by DNMT3A deficiency.

**Figure 4.**
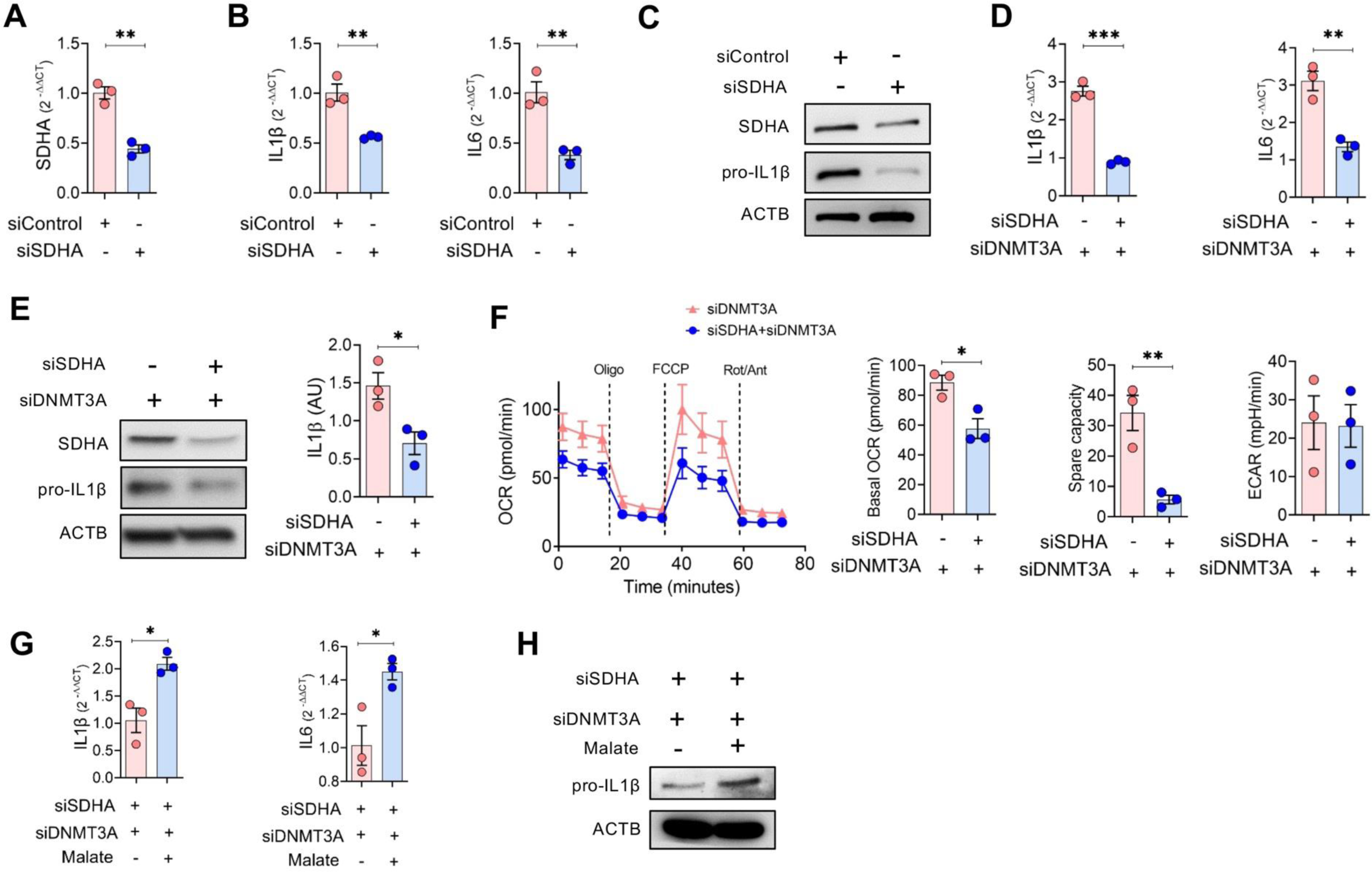
SDHA/Malate axis regulates pro-inflammatory and metabolic profile in macrophages. (**A, B**) mRNA expression of *SDHA*, *IL1β* and *IL6* and in PBMC-derived macrophages from HF patients with *DNMT3A* CHIP mutations after transfection with siControl and siSDHA for 48 hours (n=3). (**C**) SDHA and pro-IL1β protein level in PBMC-derived macrophages from HF patients with *DNMT3A* CHIP mutations after transfection with siControl and siSDHA for 48 hours (n=2). (**D**) *IL1β* and *IL6* mRNA level in DKD *SDHA* and *DNMT3A* PBMC-derived human macrophages (n=3). (**E**) Pro-IL1β protein followed by the densitometric quantification of relative pro-IL1β protein expression in double knock down (DKD) *SDHA* and *DNMT3A* PBMC-derived human macrophages (n=3). (**F**) OCR measurement in DKD *SDHA* and *DNMT3A* PBMC-derived macrophages (n=3). Data are shown as OCR graphs (left) representative of three independent experiments and summary bar graphs of basal respiration, spare capacity and ECAR (right) (n=3). (**G**) *IL1β* and *IL6* mRNA level in DKD *SDHA* and *DNMT3A* PBMC-derived human macrophages followed by malate (5mM) treatment for further 24 hours (n=3). (**H**) Pro-IL1β protein in DKD *SDHA* and *DNMT3A* PBMC-derived human macrophages followed by malate 5mM treatment for further 24 hours (n=2). Statistical significance was assessed by a two-tailed unpaired Student’s *t*-test and. *p <0.05, **p <0.01, ***p< 0.001.

### DNMT3A deficiency induced malate production stimulates a pro-inflammatory phenotype in a paracrine manner

Apart from the intrinsic pro-inflammatory and metabolic effects of DNMT3A in macrophages, the paracrine functions of DNMT3A-associated macrophages on non-mutated macrophages may contribute to the general pro-inflammatory phenotype in patients with DNMT3A mutations, which generally carry only a relatively small percentage of mutant cells. To determine the paracrine activity of DNMT3A-deficient macrophages, we applied conditional media (CM) to naïve macrophages and evaluated the inflammatory phenotype and mitochondria activity. The CM of siDNMT3A macrophages induced a pro-inflammatory response in naïve macrophages in comparison to CM of siControl macrophages (Figures 5A and 5B). This pro-inflammatory phenotype is concomitant with enhancement of OCR level and mitochondria spare capacity in naïve macrophages which are treated with CM of siDNMT3A macrophages (Figure 5C). Interestingly, when we applied the CM of the double knock down *SDHA* and *DNMT3A* macrophages to naïve macrophages, both the pro-inflammatory response and mitochondria OCR are reduced (Figures 5D and 5F) which revealed a functional role of SDHA in the paracrine pro-inflammatory activation. Given that malate is known to elicit pro-inflammatory responses, we determined whether the exogenous addition of malate on the macrophages could mimic the paracrine inflammatory activation. Indeed, malate pretreatment augmented LPS-induced expression of *IL1β* and *IL6* mRNA (Figure 5G) also *IL1β* protein (Figure 5H) which concomitant with increased OCR and spare capacity (Figure 5I). Altogether, these data show DNMT3A deficiency not only affects cell-intrinsic immunometabolic effects but also can induce pro-inflammatory activation associated with increased mitochondria oxygen consumption in wild type macrophages via the SDHA/malate axis in a paracrine manner.

**Figure 5.**
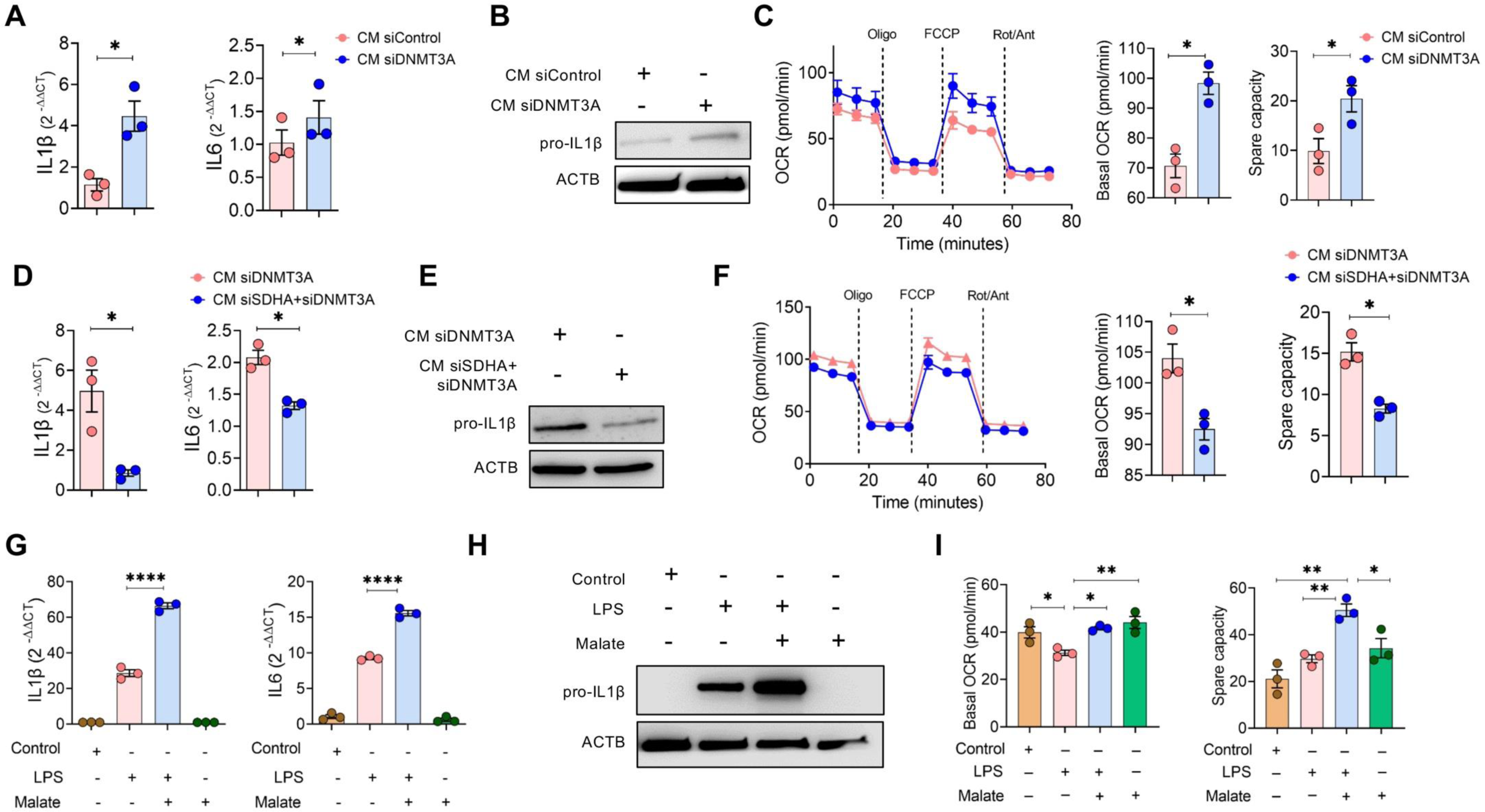
DNMT3A CHIP macrophages induces pro-inflammatory phenotype in paracrine manner. (**A**) mRNA level of inflammatory cytokines, *IL1β* and *IL6* in PBMC-derived macrophages from healthy donors that were subjected to CM of siDNMT3A and siControl macrophages for 72 hours (n=3) (**B**) Pro-IL1β protein in PBMC-derived macrophages from healthy donors that were subjected to CM of siDNMT3A and siControl macrophages for 72 hours (n=2). **(C)** OCR measurements of human PBMC-derived macrophages (*n* = 3) which are treated with CM of siDNMT3A and siControl macrophages for 48hrs hours. Data are shown as OCR graphs (left) representative of three independent experiments, and summary bar graphs of basal OCR and spare capacity (right). (**D-E**) mRNA level of inflammatory cytokines, *IL1β, IL6* (n=3) and pro-IL1β protein (n=2) in PBMC-derived macrophages from healthy donors that were subjected to CM of DKD of siSDHA and siDNMT3A macrophages for 72 hours, respectively (**F**) OCR measurements of human PBMC-derived macrophages (*n* = 3) which are treated with CM of DKD of siSDHA and siDNMT3A macrophages for 48hrs hours. Data are shown as OCR graphs (left) representative of three independent experiments, and summary bar graphs of basal OCR and spare capacity (right). **(G-H)** mRNA level of inflammatory cytokines, *IL1β, IL6* and pro-IL1β protein in PBMC-derived macrophages from healthy donors which were pretreated with malate (5mM) for 3 hours followed by LPS stimulation (100ng/ml) for further 48 hours (n=3). (**I**) OCR measurements of human PBMC-derived macrophages (*n* = 3) which are pretreated with malate (5mM) for 3 hours followed stimulation with LPS for 48 hours. Data are shown as summary bar graphs of basal OCR and spare capacity (n=3). Statistical significance was assessed by two-tailed paired Student’s *t*-test and one-way ANOVA Tukey’s multiple comparison test. *p <0.05, **p <0.01, ****p< 0.0001.

### Inhibition of SDHA reduces inflammation and improves cardiac function in DNMT3A-R882H mice

We discovered that SDHA is modulated by *DNMT3A* CHIP mutations and in turn serves as a major modulator of inflammation. Thus, we evaluated whether the pharmacological inhibition of SDH via the cell-permeable molecule dimethyl malonate ^29^ could reverse the pro-inflammatory macrophage phenotype in DNMT3A CHIP heart failure patients.

First, we determined the effect of dimethyl malonate on the inflammatory phenotype of LPS-stimulated, DNMT3A CHIP macrophages from heart failure patients and observed that dimethyl malonate treatment significantly reduced the expression of inflammatory genes (Figure 6A; Figure S4A). Strikingly, the effect of dimethyl malonate on *IL1β* expression was much more prominent in macrophages derived from DNMT3A CHIP heart failure patients (Figure 6B), suggesting the higher dependency of DNMT3A CHIP macrophages on complex II activity. These in vitro findings suggest that SDHA inhibition might be a therapeutic strategy to reduce inflammation in heart failure patients with *DNMT3A* mutations.

**Figure 6.**
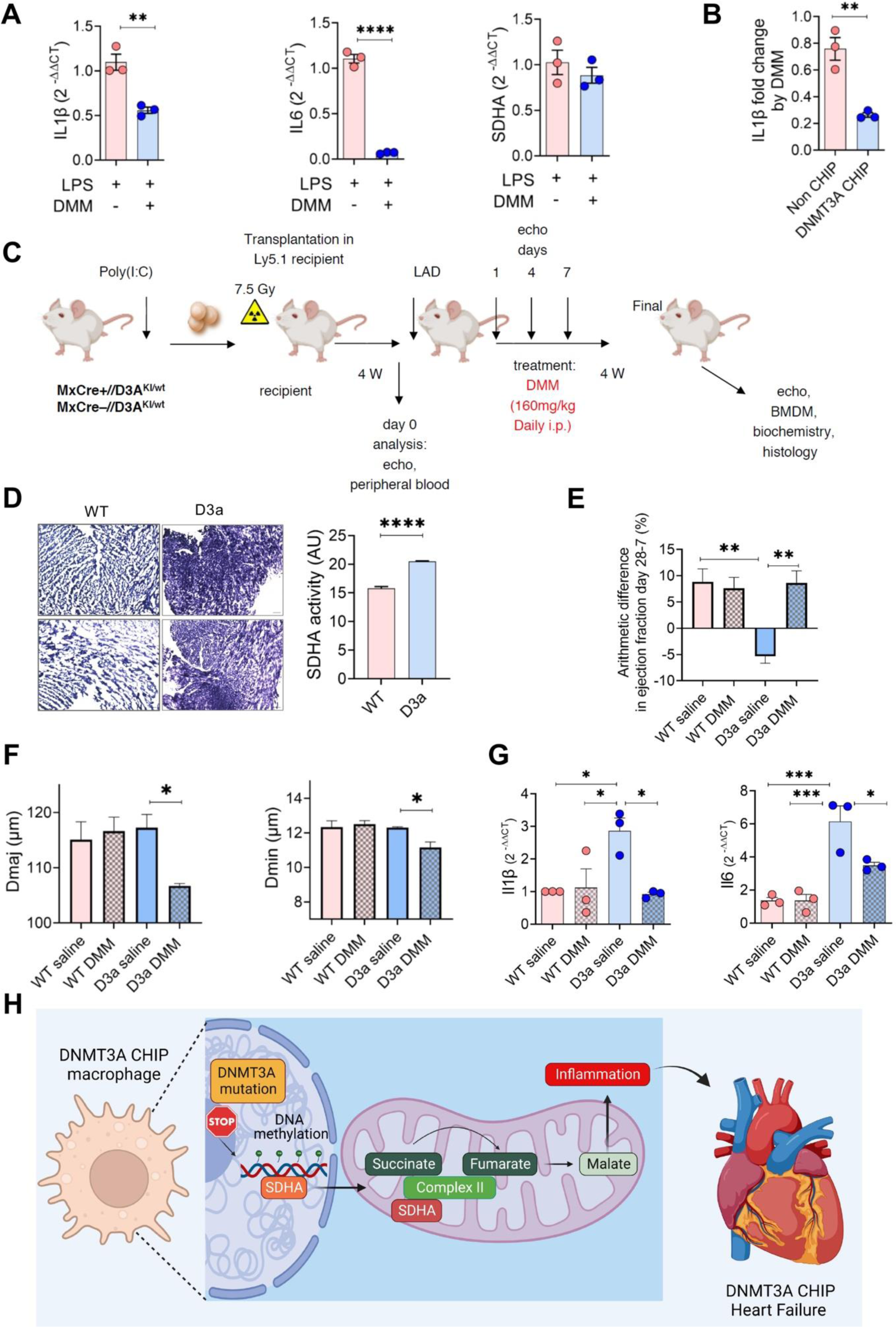
Inhibition of SDHA with dimethyl malonate improves cardiac function and reduces pro-inflammatory phenotype after myocardial infarction in *DNMT3A*-R882H mice. (**A**) mRNA level of inflammatory cytokines, *IL1β, IL6* and *SDHA* in PBMC-derived macrophages from CHIP HF patients with *DNMT3A* mutations which were pretreated with dimethyl malonate (10mM) for 3 hours followed by LPS stimulation (100ng/ml) for further 48 hours (n=3). (**B**) Fold change of *IL1β* mRNA levels between PBMC-derived macrophages from HF patients with *DNMT3A* mutations and non-CHIP after dimethyl malonate treatment (10mM) under LPS stimulation (100ng/ml) for 48 hours (n=3). (**C**) Schematic representation of ligation of the left anterior descending artery (LAD) in humanized *DNMT3A*-R882H (D3a) mouse following with dimethyl malonate treatment. (**D**) *in situ* SDHA activity in heart of transplanted mice with *DNMT3A*-mutated and WT HSCs followed by quantification analysis with Image J. (**E**) Arithmetic difference, as cardiac function indicator, in D3a mice and WT upon LAD followed by dimethyl malonate treatment. (**F**) Cardiac hypertrophy phenotype which is indicated with cardiomyocyte length (Dmaj) and diameter (Dmin) in D3a mice and WT upon LAD followed by dimethyl malonate treatment. (**G**) mRNA expression of pro-inflammatory markers including *Il1β* and *Il6* of bone-marrow-derived macrophages (BMDMs) from WT and D3a mice upon ligation of LAD followed by dimethyl malonate treatment.(H) *DNMT3A* mutations in CHIP macrophages leads to hypomethylation of *SDHA* gene which results in higher gene expression and protein level of SDHA also enhancement activity of complex II in ETC. SDHA/complexII converts succinate to fumarate followed by higher malate production in DNMT3A CHIP macrophages. Malate can augment the pro-inflmmatory response in patients with *DNMT3A* mutations which finally can lead to HF in this group of patients.Statistical significance was assessed by a two-tailed unpaired Student’s *t*-test and one-way ANOVA Tukey’s multiple comparison test. *p <0.05, **p <0.01, ***p< 0.001, ****p< 0.0001.

To assess the impact of dimethyl malonate in vivo, we used a murine model that conditionally expresses human *DNMT3A* cDNA carrying the R882H mutation ^21^. Ly5.1 mice (C57BL/6 congenic strain that carries the differential Ptprca/CD45.1 pan leukocyte marker) were transplanted with *Mx1*-*Cre*^-^:*DNMT3A*^WT/R882H^ or *Mx1*- *Cre*^+^:*DNMT3A*^WT/R882H^ bone marrow cells before subjection to left anterior descending artery (LAD) ligation that induces myocardial infarction (Figure 6C). 4 days after induction of myocardial infarction, mice were treated daily with dimethyl malonate for seven days (Figure 6C). Mice transplanted with *DNMT3A*-mutated cells had worse cardiac function and higher cardiac SDHA activity (Figure 6D) than animals treated with wild-type cells. However, dimethyl malonate improved cardiac function as evidenced by an improved in arithmetic difference in ejection fraction (Figure 6E). Interestingly, cardiac hypertrophy as measured by cardiomyocyte length (Dmaj) and diameter (Dmin) was specifically reduced in dimethyl malonate treated mice bearing *DNMT3A*^WT/R882H^ cells (Figure 6F). Monocyte/macrophages recruitment and activation exert the regulatory roles in cardiac remodeling and fibrosis^30^. To explore the monocyte/macrophage phenotype in mice transplanted with *DNMT3A*-mutated cells followed by LAD ligation, we checked the inflammatory phenotype of bone-marrow derived macrophages (BMDMs). Dimethyl malonate treatment decreased the expression of pro-inflammatory markers including *Il1β* and *Il6* especially in mice transplanted with *DNMT3A*-mutated cells followed by LAD ligation (Figure 6G). These data manifest that the induction of *DNMT3A* mutation in hematopoietic cells aggravates cardiac dysfunction after myocardial infarction. Inhibition of SDH by dimethyl malonate reduced cardiac inflammation and improved cardiac function specifically in mice bearing DNMT3A-mutant cells (Figure 6H).

## Discussion

Probing the unexplored metabolic aspects in CHIP-associated CVD, our study shows that mitochondrial metabolism through the oxidation of succinate following complex II/SDHA function is one of the central hubs to shape the inflammatory phenotype of macrophages with *DNMT3A* CHIP mutations or DNMT3A deficiency. Our major findings include (i) activation of complex II in macrophages isolated from DNMT3A CHIP mutations and DNMT3A deficient macrophages; (ii) hypomethylation and upregulation of SDHA in DNMT3A CHIP heart failure patients; (iii) Enrichment of malate, a downstream metabolite of SDHA, in serum from DNMT3A CHIP heart failure patients and induces a pro-inflammatory phenotype in macrophages; (iv) genetic or pharmacological inhibition of SDH decreases the pro-inflammatory phenotype in macrophages with *DNMT3A* CHIP mutations and DNMT3A-deficient macrophages; and (iv) the metabolic intervention with the SDH inhibitor dimethyl malonate ameliorated the inflammatory response and improved the cardiac function in a heart failure animal model, which harbor myeloid-specific *DNMT3A* mutation.

We identified that mitochondrial metabolism majorly complex II/SDH as a key regulator of the inflammation in CHIP associated pathology. Beyond the potential role of mitochondrial metabolism in driving inflammatory functions, mitochondrial respiration is required for HSC proliferation ^31^. Higher mitochondria respiration in DNMT3A CHIP cells could explain why this specific CHIP mutation is the most prevalent clone in humans^32^. However, longitudinal studies are warranted to confirm whether the mitochondria metabolite fluctuations correlate with CHIP clone size.

The role of complex II/SDH in driving inflammation has previously been shown in different pathological contexts. For example, during ischemia-reperfusion injury in the heart and brain, reverse electron transport in complex I followed by succinate oxidation in complex II/SDH leads to ROS production ^27,33^. Recently, higher activity of complex II/SDH as a consequence of SDHA gain-of-function mutation was reported in patients with persistent polyclonal B cell lymphocytosis ^34^. Similarly, our study revealed that DNMT3A CHIP-associated heart failure also seems to be driven by complex II/SDH hyperactivity and accompanying metabolic deregulation that was reflected in higher OCR levels in macrophages with *DNMT3A* CHIP mutations as well as in DNMT3A-deficient macrophages. Although increased mitochondrial activity is often associated with increased ROS production, this was not observed in DNMT3A silenced macrophages. The latter finding is consistent with findings from another group ^35^.

The increased SDHA in CHIP carrying heart failure patients is consistent with the marked elevation in its downstream product, malate. In addition, our data further suggest malate as a driver of the inflammatory phenotype and cardiac dysfunction in heart failure patients carrying *DNMT3A* CHIP mutations. The accumulation of malate during the pro-inflammatory response has been reported in macrophages ^27,36^ and associated with a higher risk of arterial fibrillation and heart failure in individuals with a high cardiovascular risk ^37^. Increased serum levels of malate and its potential role in driving inflammation, may explain how a relatively low number of *DNMT3A* mutant cells can support a state of chronic, low-grade inflammation in patients with *DNMT3A* CHIP mutations ^32^. Moreover, the malate derived from DNMT3A mutant macrophages may also exhibit effects on the structural cells of the heart. Mechanistically, this would likely involve malate-driven epigenetic modifications, or be mediated indirectly by metabolites downstream of malate e.g. fumarate and 2HG, that also act as epigenetic modifiers ^38,39^.

DNMT3A catalyzes the *de novo* methylation of cytosine residues in DNA. Previously, we ^8^ and others ^23^ reported DNA hypomethylation in blood samples of COPD and CVD patients with *DNMT3A* CHIP mutations. Fitting with this, our analyses indicate that *SDHA* was also upregulated in macrophages carrying *DNMT3A* CHIP mutations. This observation implies a clear link between *SDHA* DNA methylation and the transcriptional regulation of SDHA. While it is clear that DNMT3A exerts a number of different functions during inflammation, e.g. the type I IFN response ^35^ and antiviral innate immunity ^40^, our data point to a critical role of DNMT3A-dependent changes in DNA methylation in the control of immune cell metabolism.

Importantly, based on the immunometabolic characteristics of patients with *DNMT3A* mutation, metabolic modulation might provide a potential therapeutic approach. In corroboration, intravenous infusion of SDH inhibitor – dimethyl malonate, in a mouse model that mimics DNMT3A-mutated CHIP that has been subjected to myocardial infarction demonstrated a significant improvement of cardiac function, and reduction of cardiac inflammation. Although dimethyl malonate is not tested in any clinical trials, the natural SDHA inhibiting compounds with reduced toxicity such as itaconate and its derivatives ^41^ can be considered as promising approaches for future clinical trials. Alternatively, targeting inflammation with antibodies directed against IL-1β and IL-6, or NLRP3 or STAT-3 inhibitors may mitigate the effects of CHIP, both by reducing clonal expansion and CHIP-dependent inflammation. Certainly, IL-1β neutralization in the Canankinumab Anti-inflammatory Thrombosis Outcomes (CANTOS) trial suggested that TET2 variants may respond better to canakinumab than those without CHIP ^42^. An added advantage of metabolic modulation includes manipulation of the signaling pathways that are necessary and upstream of the inflammatory response.

In conclusion, our findings revealed SDHA as a disease modifier in DNMT3A CHIP heart failure, driving malate accumulation and inflammatory reprogramming of macrophages. This study reports for the first time how the metabolic regulatory functions can be involved in DNMT3A CHIP pro-inflammatory and cardiac phenotypes which can be used as a promising individualized therapeutic approach in CVD patients with *DNMT3A* CHIP mutations.

## Acknowledgments

We would like to acknowledge excellent technical contributions of Laura Kahnke and Katharina Leib. Funding: This study was funded by the DFG SFB-1213 (Project A01, A05 grants to SSP, A10N grant to RS) and ERC Consolidator Grant (866051 to SSP). RS, SD, SSP are supported by Excellence Cluster CPI (Exc2026).

## Author Contributions

SM, RS, SD, SSP designed the research study. SM, IH, GZ, SK, XL, MS, SG, MP conducted the experiments and acquired the data. SM, SK, CMT, MS, MR, IF, RJR, ML, RS, SD, SSP analyzed the data. FB, MHS, ML provided bioinformatic support. SC, KK, SK, CFV, RB, AZ, WS provided clinical samples and CHIP mutational status. SM, RS, SD, SSP took the lead in writing the manuscript. IF, AZ and WS provided critical feedback and helped shape the research, analysis and manuscript.

## STAR Methods

### Study population

For COPD patients, we used the COSYCONET as a German multicenter prospective observational trial, which recruited 2741 patients aged 40 years and older with diagnosis of COPD between 2010 and 2013 in 31 study centers. COSYCONET was approved by the ethics committees at all participating sites, and all participants provided written consent. The study is registered at ClinicalTrials.gov (NCT01245933). The Study protocol V1.6 from the 23.05.2011 states full compliance with national laws, ICH Guideline for Good Clinical Practice (GCP) E6 from June 1996 and the declaration of Helsinki. The study subjects of COSYCONET gave written consent upon study inclusion for genetic analysis of blood samples. Participants also gave written consent, that results genetic analysis would not be reported to them. All blood samples for CHIP sequencing were drawn at visit 1 (between 2011 and 2013). We estimate the risk of malignancy with 0.5-1% per year in CHIP positive patients. DNA sequencing and data analysis took place 6-8 years after sampling, the study team considered this no longer relevant for the patient’s safety. For HF patients, and healthy controls, whole blood samples were taken at outpatient clinic visits patients and sequenced for the presence of DNMT3A CHIP-driver mutations. Sequencing was done as described previously^43^. In total, blood samples of 11 patients with heart failure and DNMT3A driver mutations and 13 patients with heart failure without CH mutations were used for this study. In addition, six patients without heart failure were used as controls. The obtaining of the blood samples was in the frame of the UCT-Project-Nr.: KardioBMB#2022-001 with Title: ‘Epigenetic and metabolic contribution to inflammatory phenotypes of CHIP’ and KardioBMB#2020-004: ‘Clonal hematopoiesis in heart failure patients; Amendment 1: Metabolic status and the connection with DNMT3A CHIP in patients with HFrEF’. Informed consent was obtained from all patients. The study was approved by an institutional review committee of the University Hospital of the Johann Wolfgang Goethe University in compliance with internal standards of the German government, and procedures followed were in accordance with institutional guidelines and the Declaration of Helsinki

### DNA methylation analysis

DNA was isolated from the EDTA blood samples obtained from the patients using the QIAamp DNA mini Kit (Qiagen), according to the manufactures protocol. Qubit dsDNA HS Assay kit was used to measure the DNA concentration (Q32851, Invitrogen). For DNA methylation analysis, we applied 200–500 ng of DNA as input. The Infinium Human-Methylation EPIC BeadChip (850k) (WG-317, Illumina) was used to determine the DNA methylation status of more than 850.000 CpG sites, respectively following the producer’s guidelines (1). On-chip quality metrics of all samples were carefully controlled. The RnBeads software (version 2.0) was used for analysis of the DNA methylation arrays ^44^.

Raw intensities derived from IDAT files passed quality control on probes and samples. RNbeads was run on the IDAT files to remove sex chromosomes and SNPs to avoid confounding factors. We used the following commands for this as part of the *rnb.options* function: min.group.size=1, differential.enrichment = TRUE, import.table.separator = ",", normalization = NULL, normalization.method = "illumina", normalization.background.method = "methylumi.noob", filtering.snp = "yes", filtering.sex.chromosomes.removal = TRUE, filtering.missing.value.quantile = 0, exploratory.columns = SampleID, differential.comparison.columns =GroupID, differential.comparison.columns.all.pairwise = GroupID. We used the differential methylation analysis capabilities to contrast samples with and without observed CHIP mutation for 26 COPD patients and 8 patients with CHF. Individual site-based methylation levels were calculated and subsequently used for region-based methylation level assessment in promoters of genes, by considering the mean methylation of all CpG sites on the array that belonged to the gene promoter.

We used an MA-plot to illustrate the differences in promoter methylation between CHIP and no CHIP patients using a differential methylation *p*-value cutoff of 0.01 and methylation fold change of at least 0.2 to remove genes with marginal differences. We highlighted the top 10 genes with the strongest DNA methylation differences.

### Sample Preparation for NGS and High-Throughput Sequencing

Before sequencing, the pooled libraries were diluted and denatured according to the NextSeq System Denature and Dilute Libraries Guide (Illumina) and 1% PhiX DNA was added. The pooled libraries were sequenced on a NextSeq 500 sequencer (Illumina) using the NextSeq 500/550 Mid Output, version 2 kit (300 cycles) according to the manufacturer’s instructions. Briefly, the sequencer was operated in a paired-end sequencing mode with 2 × 150 bp read length and 2 × 8 bp index read length. The BCL files were demultiplexed and converted to FASTQ files using the FASTQ Generation tool on BaseSpace (Illumina). The median coverage across all BMC samples was 4282× before UMI family clustering and 630× with inclusion of UMIs.

Variant Calling and Annotation Strategies: Read quality was assessed using FastQC. FASTQ files from each patient were merged and reads were grouped into unique molecular identifier (UMI) families using the UMI-tools software package ^45^. Reads were mapped to the hg19 draft of the human genome using Burrows-Wheeler Alignment–MEM ^46^. The ‘dedup’ command of the UMI-tools software package was used to remove polymerase chain reaction duplicates with the same UMIs and alignment coordinates. Variant calling was performed using FreeBayes without allele frequency threshold, a minimum alternate read count of 2, and a minimum base and mapping quality of 20. Variant effect prediction and variant annotation was performed using SnpEff and SnpSift (2,3,4).

The identified variants were processed and filtered using the R programming language, version 3.3.1 (R Programming). Common single-nucleotide polymorphisms with a minor allele frequency of at least 5% in either the 1000 Genome Project, Exome Variant Server, or ExAC databases were excluded ^47^. In addition, variants with a low mapping quality (<20) and variants occurring in 8% or more of the patients in the studied cohort were considered as technical artifacts and excluded. Furthermore, variants covered with fewer than 100 reads in at least 1 set of amplicons (CAT A or CAT B), variants called with only 1 of the set of amplicons (CAT A or CAT B), single-nucleotide polymorphisms identified as common in the single-nucleotide polymorphism database (≥1% in the human population), and variants with sequence ontology terms “LOW” or “MODIFIER” were filtered out. According to previously established definitions, all variants with a variant allele fraction (VAF) of at least 0.02 (2%) were considered; VAF was calculated by using the formula VAF = alternate reads / (reference + alternate reads). Variants with a VAF of 0.45 to 0.55 were not considered to exclude potential germline variants. The variants were further validated on the basis of being reported in the literature and/or the Catalogue of Somatic Mutations in Cancer ^48^ and ClinVar ^49^.

### Metabolomic and cytokine analysis

Plasma samples were isolated at study visits in the according study center. For Isolation and direct processing of samples the studies (SOP) was followed. Blood vials were centrifuged and frozen within 1h after sampling. A temperature log protocol was followed. All biomaterials were stored in the central biobank. The transfer of the samples from the study centers to the central biobank and all related procedures are defined by SOPs. The transportation system is established, and every step described in the corresponding SOP. The samples are transported under dry ice pellets. After receipt at the biobank the samples are inspected immediately, imported in the database and stored at -80°C until further processing.

Plasma samples were extracted in ice-cold 85% methanol (10 µL/µL), shortly vortexed and lysed with one freeze-thaw cycle. The homogenate was centrifuged (15,000 g, 5 minutes, 4 °C). The supernatant was collected and the samples were evaporated to dryness in a Concentrator Plus (Eppendorf, Wesseling-Berzdorf, Germany). Samples were reconstituted in 50 µL water + 0.5% formic acid, transferred to autosampler vials and subsequently analyzed via liquid chromatography coupled to tandem mass spectrometry (LC-MS/MS). Negative ionization ESI-LC MS/MS was performed on an Agilent 1290 Infinity LC system (Agilent, Waldbronn, Germany) coupled to a QTrap 5500 mass spectrometer (Sciex, Darmstadt, Germany). Ion source parameters were as follows: CUR 30 psi, CAD medium, Ion Spray Voltage - 4500 V, TEM 400 °C, GS1 45 psi, GS2 25 psi. TCA metabolites were identified with authentic standards and/or via retention time, elution order from the column and 1-2 transitions. For quantification, specific MRM transitions for every compound were normalized to its appropriate standards in a standard curve. Reversed-phase LC separation was performed by using a Waters Acquity UPLC HSS T3 column (150 mm × 2.1 mm, 1.8 µm). Compounds were eluted with a flow rate of 0.4 ml/min and with the following 10 min gradient: 2% B for 1.5 min, a 3 min gradient to 100% B, a cleaning and equilibration step. Solvent A consisted of 100 % water containing 0.5 % formic acid and solvent B consisted of 100 % methanol containing 0.5 % formic acid. Column oven temperature was set to 40 °C, and the autosampler was set to 6 °C. Injection volume was 2.5 µL.

Macrophage supernatant IL6 was measured by human IL6 Quantikine ELISA Kit (R&D) based on manufacture’s instruction.

### Generation of human and mouse macrophages and cell culture

Human and mouse macrophages were generated from peripheral blood mononuclear cells (PBMCs) and bone marrow (BM) cells, respectively as previously described. Briefly, healthy donors, CHIP and non-CHIP PBMCs isolated from buffy coats obtained from the blood bank of the Universities of Giessen and Marburg Lung Center and Frankfurt Universities Hospital using Ficoll density gradient centrifugation. Platelets and red blood cells (RBC) were removed by two washing steps with RBC lysis buffer (BD Biosciences) and phosphate-buffered saline (PBS), respectively. The pellets were passed through the filter and then resuspended in ice cold Miltenyi buffer followed by adding magnetic-activated cell sorting (MACS) CD14 beads (Miltenyi Biotec) and incubation at 4°C which terminated by adding Miltenyi buffer after 15 minutes. The tubes were centrifuged at 300 x g for 5 minutes at room temperature. The supernatant was removed and the pellets were resuspended in ice cold Miltenyi buffer. The positive CD14 selection was performed by using LS 2 (Miltenyi Biotec) columns were already equilibrated by adding Miltenyi buffer. The cell solution was applied onto the columns and then washed three times with Miltenyi buffer. After the last wash, the columns were filled with Miltenyi buffer and then plunged to flushing out the CD14-positive cells which were collected with centrifugation at 300 g for 5 minutes with acceleration. The supernatant was discarded and the cell pellet was resuspended in RPMI-1640 (20 ml) (Gibco) supplemented with 10% (v/v) fetal-bovine serum (FBS; Life technology) and penicillin-streptomycin (Pen/Strep; Gibco). The cells were counted and plated at 0.5 x 10^6^ cells/ml in the media plus macrophage colony-stimulating factor (M-CSF,25 ng/ml, R&D Systems). Cells were maintained at 37°C, 5% CO_2_ for 5 days, to allow differentiation into macrophages with one media change in 3^rd^ day. Thereafter, macrophages were transfected with stealth RNAi siRNAs against DNMT3A (HSS176225, Thermofisher) and siRNA against SDHA (Catalog no.: SI00060445, Qiagen) and a scramble negative control (AllStars Neg. siRNA, Catalog no.: 1027292, Qiagen) using the HiPerFect Transfection Reagent (Qiagen) in an optimum serum-free medium based on the manufacturer’s protocol.

Regarding mouse macrophages, the tibia and femur from mice were dissected, and each bone was subsequently flushed with 15 ml of RPMI 1640 medium with 1% Pen/Strep. The cells went through a 40-μM cell strainer, centrifuged, and then resuspended in RPMI 1640 medium which contained 10% FCS, 1% Pen/Strep, and mouse M-CSF(20 ng/ml; Roche) and plated on a six-well plate. We changed the medium on alternate days with RPMI 1640 medium contained 10% FCS, 1% Pen/Strep, and mouse M-CSF (20 ng/ml) until undifferentiated macrophages were obtained. For polarization of human and mouse macrophages toward the pro-inflammatory macrophage, we stimulated them with LPS (100 ng/ml) for 48 hours.

### RNA isolation, complementary DNA synthesis, and quantitative PCR

Total RNA was extracted with RNeasy Mini Kit (Qiagen, Hilden, Germany). Then, we was transcribed RNA into complementary DNA (cDNA) using the high-capacity cDNA reverse transcription Kit (Applied Biosystems, Waltham, USA) according to the manufacturer’s instructions. Also, quantitative PCR (qPCR) was performed using SYBR Green PCR Master mix and the StepOne real-time PCR System (Applied Biosystems, Waltham, USA) at the following conditions: 10 min at 95°C, followed by 40 cycles of 30s at 95°C, 30s at 58° to 60°C. Analysis was done using the StepOne plus software and GraphPad Prism. Expression was determined using the ΔΔCT method followed by fold change calculation. CT-values were normalized to the housekeeping gene-encoding hypoxanthine-guanine phosphoribosyl transferase (*HPRT1*) using the equation ΔCT = CT_reference_ – CT_target._ The primer sequences were designed using sequence information obtained from the National Center for Biotechnology Information database and purchased from Sigma-Aldrich and shown table S1.

### Western Blotting

The cells were lysed in RIPA lysis buffer containing protease and phosphatase inhibitors followed by clearing step through high-speed centrifugation. Proteins were separated using 10% polyacrylamide gels and transferred to polyvinylidene difluoride membranes. Following the blocking with 5% bovine serum albumin, the membranes were incubated with a primary antibody overnight at 4°C. After washing with tris-buffered saline containing Tween 20 (TBST) for 3 times, the blots were incubated with secondary antibodies coupled with horse radish peroxidase diluted in 5% milk dissolved in TBST buffer. The protein-antibody conjugates were detected using an enhanced chemiluminescence detection system. Protein expression was quantified using band intensity values (in arbitrary units) that were normalized to β-actin (ACTB) by ImageJ software. The Western blots shown in the figure are representative of 3 independent experiments unless it mentioned in the figure legends. Antibody details were shown in table S2.

### Metabolic flux analysis

We used A Seahorse XFe96 extracellular flux analyzer (Seahorse Bioscience, Agilent Technologies) to measure OCRs and ECARs. Mitochondrial perturbation experiments were carried out by sequential addition of 1.5 μM oligomycin (Agilent Technologies), 1 μM FCCP (carbonyl cyanide 4-(trifluoromethoxy) phenylhydrazone; Agilent Technologies) and 0.5 μM rotenone/antimycine (Agilent Technologies). For monitoring the OCR of intact mitochondria, CD14-MACS sorted monocytes were seeded in 96 wells Seahorse plate in 15*10^4^ cells/per well and differentiated to macrophages in RPMI-1640 media (Gibco) which contained with 10% (v/v) FBS (Life technology), 1% Pen/Strep plus M-CSF(25 ng/ml, R&D Systems) for 5 days with one media change in 3^rd^ day. Thereafter, the media was removed and replaced by MAS buffer (70 mM sucrose, 220 mM mannitol, 10 mM KH_2_PO_4_, 5 mM MgCl_2_, 2 mM HEPES and 1 mM EGTA), then macrophages treated with saponin (25 μg/ml) as membrane permeabilizer (Sigma-Aldrich), which was followed by addition of 5 mM pyruvate, 2.5 mM malate, 1 mM ADP and 1 μM rotenone for monitoring complex I-driven respiration; 10 mM succinate, 1 mM ADP and 0.04 mM malonate for monitoring complex II-driven respiration; 0.5 mM duroquinol, 1 mM ADP and 0.02 mM antimycin A for monitoring complex III-driven respiration; 0.5 mM TMPD, 2 mM ascorbate, 1 mM ADP and 20 mM sodium azide (all from Sigma-Aldrich) for monitoring complex IV-driven respiration. OCR changes upon the substrate addition were calculated relative to the preinjection rate.

### SDH enzyme activity

For tissue section preparation, heart tissues were harvested, snap-frozen and stored at −80°C. The tissue blocks which were embedded in O.C.T. compound cut in 6 μm thickness at slow and constant speed and placed up on Superforst microscopy slides. Slides were stored at −80°C for further enzyme activity staining procedure. SDH enzyme activity staining on tissue sections were carried out as described by Miller et. al ^50^. Briefly, we used 0.1 M Tris-HCl buffer pH 8.0 for SDH activity. In the SDH-specific buffer, 10% polyvinyl alcohol was dissolved at 60 °C under stirring until the mixture was clear. Then, SDH-specific assay media were prepared by adding 60mM succinate, 5mM sodium azide, 0.2 mM phenazine and 5 mM nitroblue tetrazolium chloride (NBT) including negative control reactions with the addition 250 mM malonic acid (for SDH inhibition). Assay medium containing the enzyme-specific substrates was applied to cover the whole tissue section slide. Enzyme reactions were carried out at RT for around 15 min or until high staining was visible and stopped by the removal of the incubation medium and washed with warm PBS.

### MitoSOX ROS staining

Reactive oxygen species (ROS) formation in macrophages with *DNMT3A* CHIP mutations and non-CHIP was determined using the MitSOX™ red mitochondrial superoxide indicator (Invitrogen, Waltham, USA) according to the manufacturer’s instructions. PBMC-derived macrophages were grown on cover slips as explained before and treated with 5 µM MitoSOX reagent working solution for 10 min at 37 °C. Then after, the coverslips were washed three times with warm 1x PBS, and imaged using a fluorescence microscope (Keyence, Osaka, Japan).

### RNAseq analysis

Trimmomatic version 0.39 was employed to trim reads after a quality drop below a mean of Q20 in a window of 20 nucleotides and keeping only filtered reads longer than 15 nucleotides ^51^. Reads were aligned versus Ensembl human genome version hg38 (Ensembl release 104) with STAR 2.7.10a ^52^. Aligned reads were filtered to remove: duplicates with Picard 2.27.4 (Picard: A set of tools (in Java) for working with next generation sequencing data in the BAM format), multi-mapping, ribosomal, or mitochondrial reads. Gene counts were established with featureCounts 2.0.3 by aggregating reads overlapping exons on the correct strand excluding those overlapping multiple genes ^53^. The raw count matrix was normalized with DESeq2 version 1.36.0 ^54^. Contrasts were created with DESeq2 based on the raw count matrix. Genes were classified as significantly differentially expressed at average count > 5, multiple testing adjusted p-value < 0.05, and -0.585 < log2FC > 0.585. Ensembl gene annotation was enriched with UniProt data (Activities at the Universal Protein Resource (UniProt)).

### Downstream analyses

All downstream analyses are based on the normalized gene count matrix.Volcano and MA plots were produced to highlight DEG expression. A global clustering heatmap of samples was created based on the euclidean distance of regularized log transformed gene counts. Dimension reduction analyses (PCA) were performed on regularized log transformed counts using the R packages FactoMineR ^55^. DEGs were submitted to gene set overrepresentation analyses with KOBAS ^56^. The resulting bubble plot shows pathways with Benjamini-Hochberg corrected p-value < 0.05 (represented by dashed line). The larger gray circles are scaled to the number of genes comprising the respective pathway, while the smaller colored circles represent subsets found to be DEGs.

### Animal model and heart functional parameters

Mx-Cre+/Dnmt3a (Ly5.2) and Mx-Cre-Dnmt3a (Ly5.2) have been previously described^21^. B6.SJL (Ly5.1) mice were obtained from The Jackson Laboratory. Animals were housed under standard laboratory conditions. Age matched Mx-Cre+/Dnmt3a and Mx-Cre-/Dnmt3a donor mice (Ly5.2) were treated with pI:pC (400μg/mouse intraperitoneally [i.p.]) for 3 nonconsecutive days (Amersham). Donor bone marrow cells (2x106 cells) were intravenously injected into lethally irradiated (7.5Gy total body irradiation) B6.SJL recipients (Ly5.1). Recipient mice were maintained on antibiotic-containing drinking water (ciprofloxacin 50 mg/kg) 5 days pre lethally irradiation and 2 weeks post irradiation and transplantation. Peripheral blood engraftment was assessed by flow cytometry 6 weeks after reconstitution.

### Left anterior descending artery (LAD) ligation in mice

Six weeks after reconstitution myocardial infarction was induced by permanent ligation of the left anterior descending artery in female mice as described previously^57^. In brief, anaesthesia was induced with isoflurane (4%/800 ml O2/min) and maintained by endotracheal ventilation (2–3%/800 ml O2/min). Thoracotomy was performed in the fourth left intercostal space. The left ventricle was exposed, and the left coronary artery was permanently occluded. Chest and skin were closed, and anesthesia was terminated. Animals were extubated when breathing was restored. Initial myocardial injury was evaluated by measuring cardiac troponin T levels in plasma 24 h after induction of myocardial infarction.

### Echocardiographic measurements

Cardiac function in mice was investigated non-invasively on a VisualSonics Vevo2100. Echocardiographic measurements were performed without anesthesia. Systolic function was evaluated in the M-mode parasternal short-axis, B-mode parasternal long- and short-axis view. The investigator was blinded towards the study group.

### Histopathological analysis

For paraffin sections, heart tissue was rinsed in PBS, fixed in 10% buffered formalin at 4°C, dehydrated, paraffinized and sectioned on a microtome (Leitz, 5µm). Tissue sections were then stained with Hematoxylin and Eosin (MilliporeSigma) and Masson Trichrome (MilliporeSigma) according to the manufacturer’s instructions. ImageJ software (NIH) was used to quantify the cardiomyocyte diameter, minor cell diameter (Dmin, µm) and major cell diameter (Dmaj, µm). Criteria for myocyteś selection were lucid cell membrane and visible nuclei.

### Statistical analysis

We analyzed all data using Prism 6.0 and 9.0 (GraphPad Software). Regarding statistical comparisons, we used unpaired sample student’s t test for HF patients and healthy donors. For paracrine experiment in healthy donors, we used paired sample student’s t test. For comparisons of groups greater than two, we performed one-way analysis of variance followed by Tukey’s post hoc test. Data were represented as mean ± SEM.

### Resources Tables

#### Antibodies list

| Antibody | Catalog number | Company |
| --- | --- | --- |
| Anti-β actin (ACTB) | ab6276 | Abcam |
| Anti-SDHA | 5839 | Cell Signaling |
| Anti-IL1β | 12703 | Cell Signaling |

#### Primers

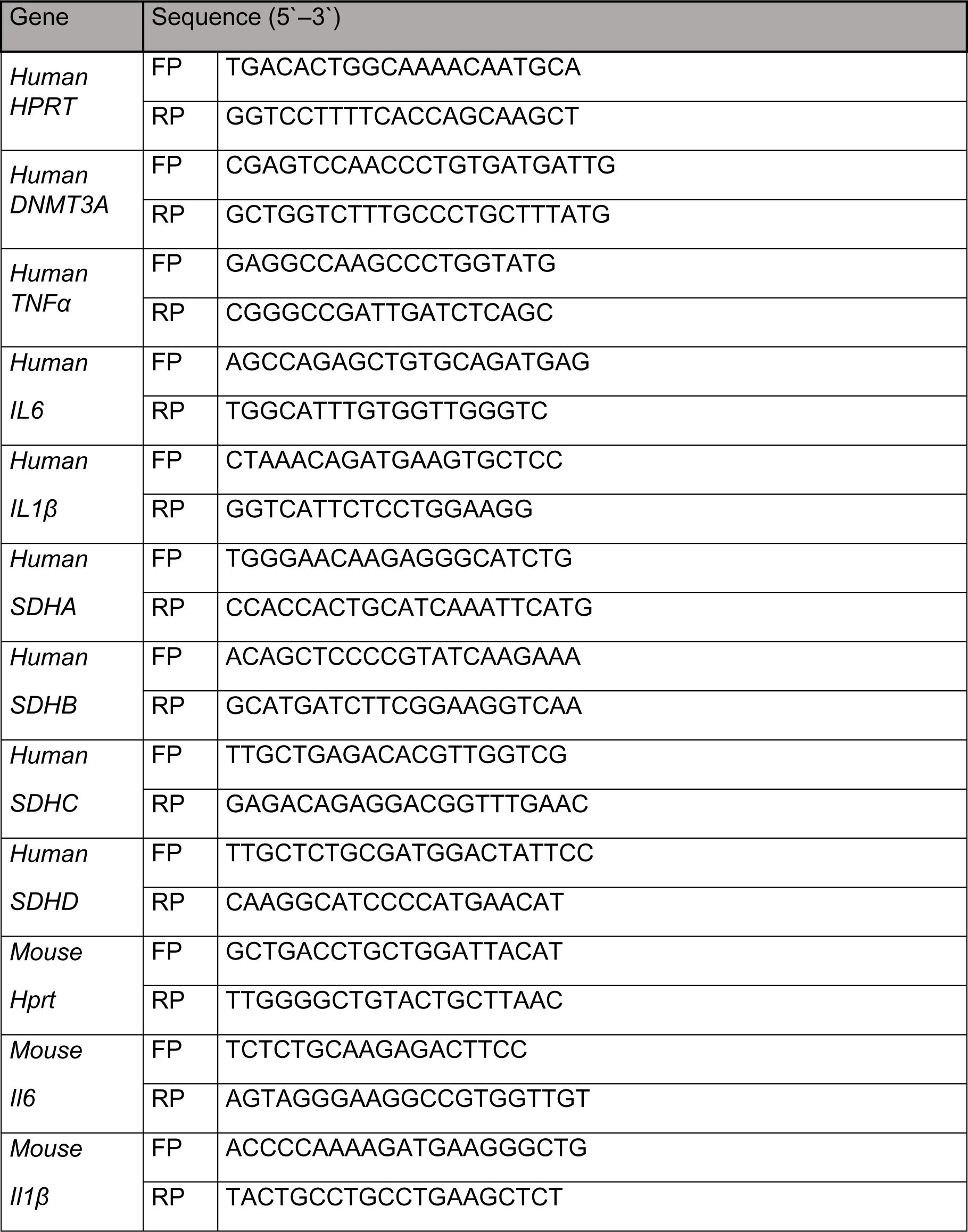

## Supplemental Information

**Figure S1.**
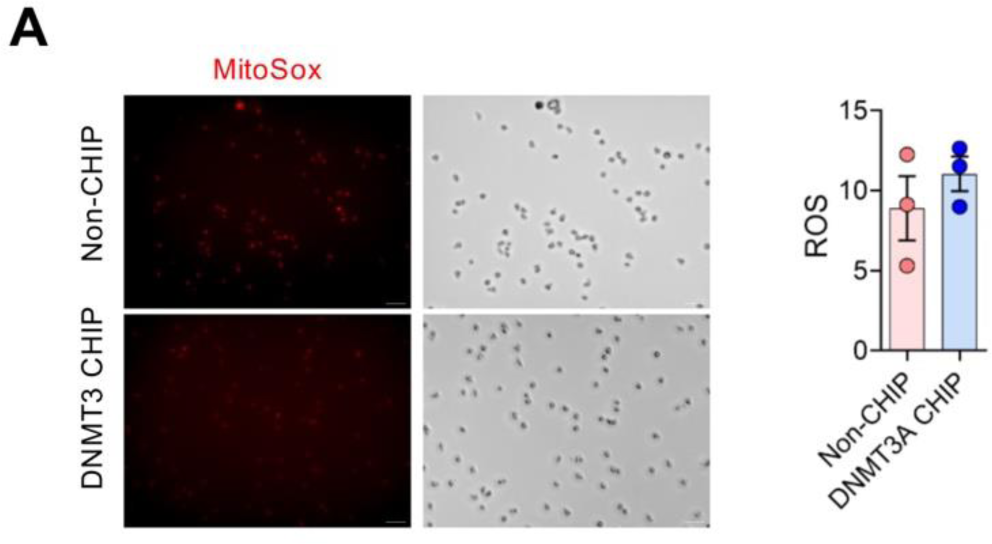
Human DNMT3A CHIP macrophages display a distinct mitochondrial metabolism with the pro-inflammatory phenotype. (**A**) Representative images (left panel) and quantification (right panel) of ROS accumulation in isolated macrophages from HF patient with *DNMT3A* CHIP mutations and non-CHIP (n = 3

**Figure S2.**
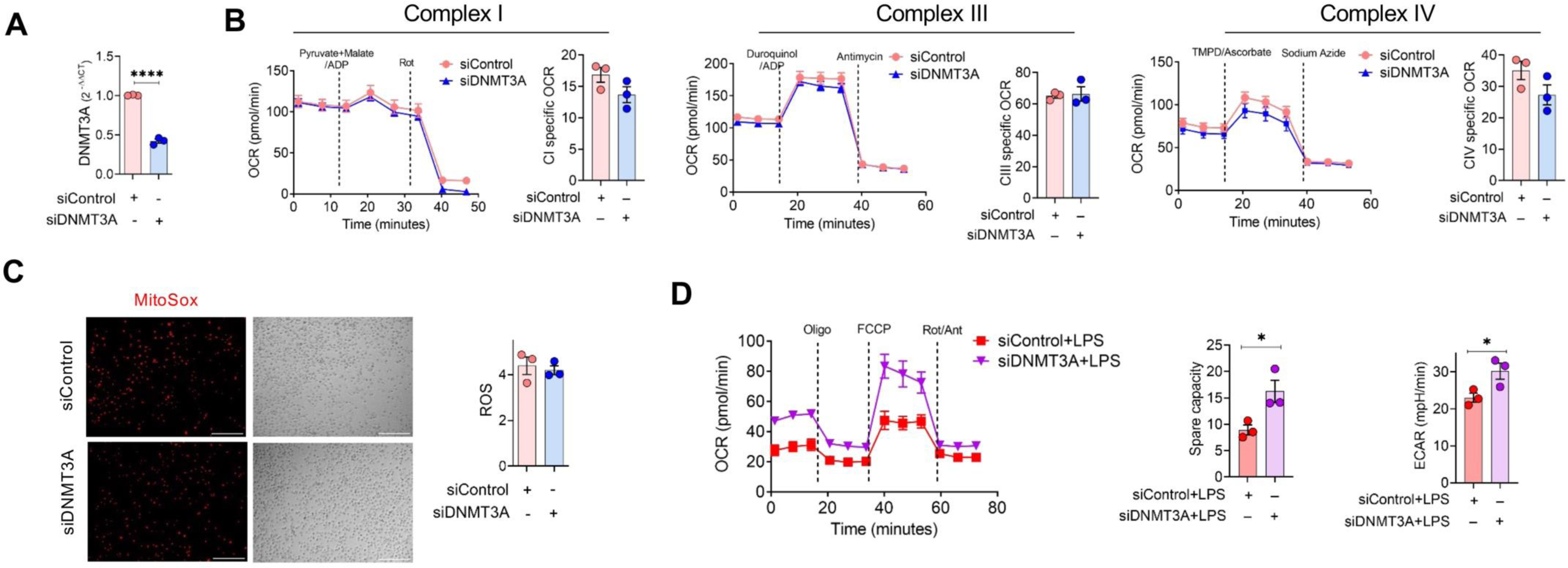
Loss of DNMT3A in macrophages induces mitochondria activity under inflammatory stimulation. (**A**) *DNMT3A* mRNA in human PBMC-derived macrophages after transfection with *DNMT3A* siRNA (n=3). (**B**) OCR measurements of permeabilized PBMC-derived macrophages after transfection with *DNMT3A* siRNA (siDNMT3A) and siControl before and after the addition of substrate and associated-complex specific inhibitors, as indicated (dashed lines) (n=3). (**C**) Representative images (left panel) and quantification (right panel) of ROS accumulation in healthy PBMC-derived macrophages after knocking down with *DNMT3A* siRNA (n=3). (**D**) OCR measurement of human PBMC-derived macrophages after transfection with siDNMT3A with LPS stimulation (100ng/ml) for 24 hours. Data are shown as OCR graphs (left) representative of three independent experiments, and summary bar graphs spare capacity and ECAR level (right) (n=3). Statistical significance was assessed by a two-tailed paired Student’s *t*-test and. *p <0.05, ****p< 0.0001.

**Figure S3.**
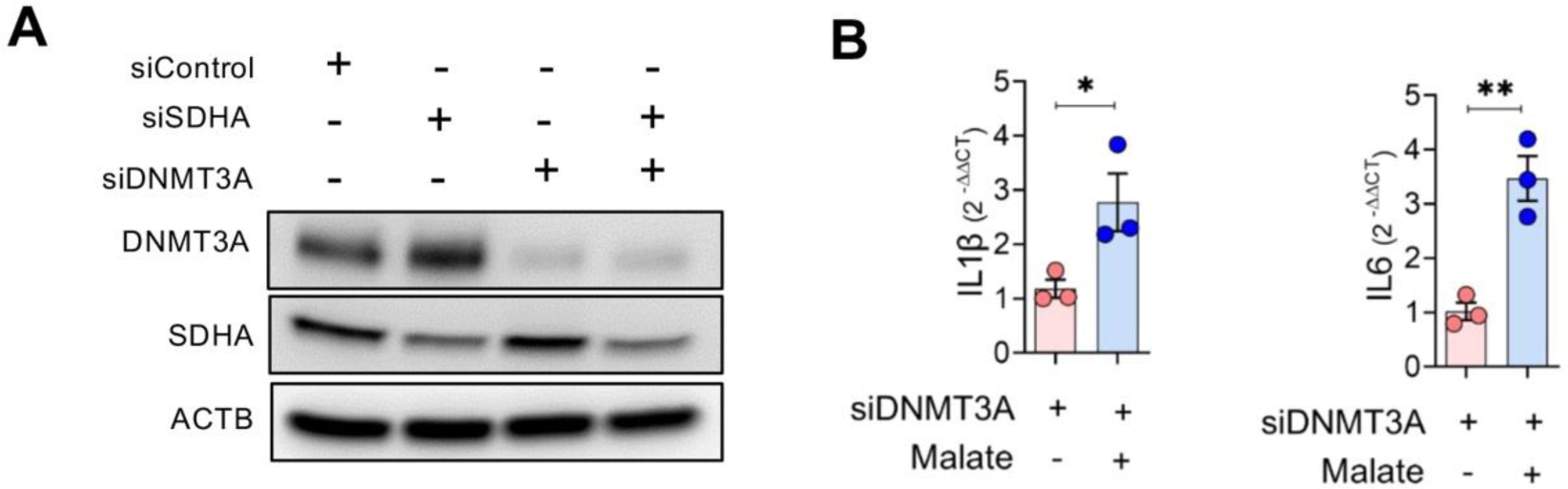
SDHA/Malate axis regulates pro-inflammatory and metabolic profile in macrophages. (**A**) DNMT3A and SDHA protein level in human PBMC-derived macrophages which are transfected with siDNMT3A and siSDHA (n=3). (**B**) mRNA expression of inflammatory cytokines including *IL1β* and *IL6* in in human PBMC-derived macrophages which are transfected with siDNMT3A followed by malate (5mM) treatment for further 24 hours (n=3). Statistical significance was assessed by a two-tailed paired Student’s *t*-test. *p <0.05, **p <0.01.

**Figure S4:**
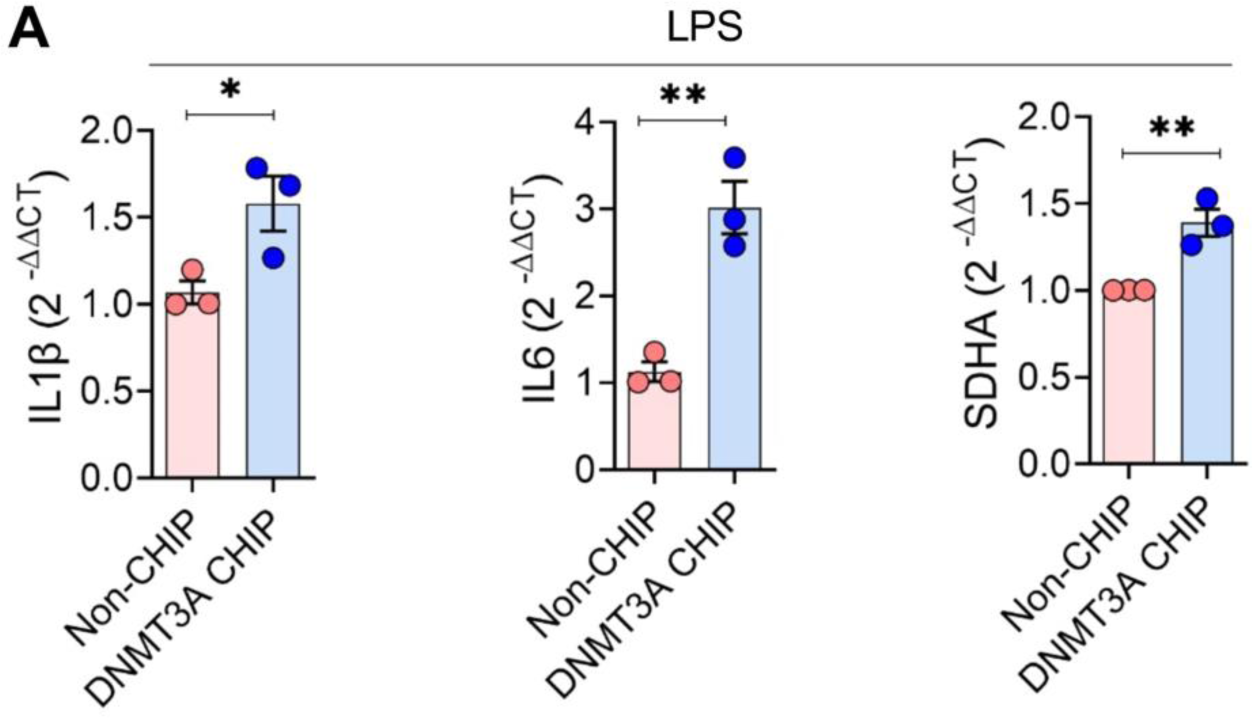
DNMT3A-associated pro-inflammatory phenotype is reduced with dimethyl malonate treatment in macrophages. (**A**) mRNA expression of inflammatory cytokines including *IL1β*, *IL6* and *SDHA* and IL1β secreted protein in PBMC-derived macrophages of HF patients with *DNMT3A* CHIP (n=3) and non-CHIP (n=3) mutations upon LPS stimulation (100ng/ml) for 48 hours. Statistical significance was assessed by a two-tailed unpaired Student’s *t*-test *p <0.05, **p <0.01.

**Table S1:**
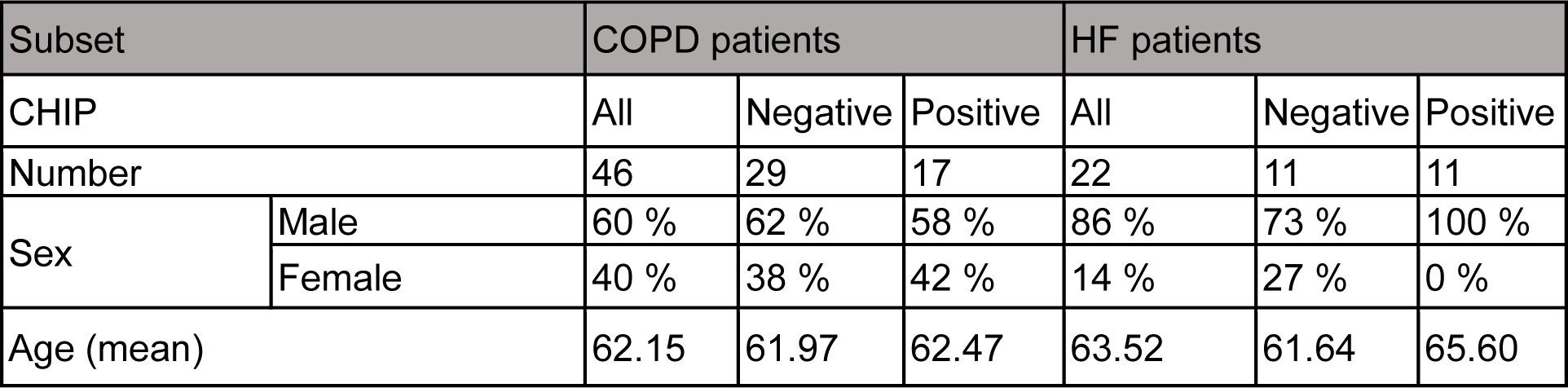
COPD and Heart failure patients’ characteristics.

| Subset |  | COPD patients |  |  | HF patients |  |  |
| --- | --- | --- | --- | --- | --- | --- | --- |
| CHIP |  | All | Negative | Positive | All | Negative | Positive |
| Number |  | 46 | 29 | 17 | 22 | 11 | 11 |
| Sex | Male | 60 % | 62 % | 58 % | 86 % | 73 % | 100 % |
|  | Female | 40 % | 38 % | 42 % | 14 % | 27 % | 0 % |
| Age (mean) |  | 62.15 | 61.97 | 62.47 | 63.52 | 61.64 | 65.60 |

**Table S2:** List of CHIP-associated somatic DNMT3A mutations identified in COPD and HF patients.

|  |  |  |  |  |
| --- | --- | --- | --- | --- |
| c.1667+1G>T<sup>#</sup> | c.2204A>G<sup>#</sup> | c.2695C>T<sup>#</sup> | c.1988C>T<sup>#</sup> | c.2007delC<sup>#</sup> |
| c.1523delT<sup>#</sup> | c.1123-1G>T<sup>#</sup> | c.2194T>C<sup>#</sup> | c.866delG<sup>#</sup> | c.2162dupA<sup>#</sup> |
| c.1135C>T<sup>#</sup> | c.886G>C<sup>#</sup> | c.892G>A<sup>#</sup> | c.2478+2T>G<sup>#</sup> | c.958C>T<sup>#</sup> |
| c.2116G>A<sup>#</sup> | c.2074C>T<sup>#</sup> | c.5627C>T<sup>#</sup> | c.2645G>T<sup>#</sup> |  |
| c.1154del<sup>*</sup> | c.1180G>T<sup>*</sup> | c.2251T>G<sup>*</sup> | c.2130_2142del<sup>*</sup> | c.1936+1G>A<sup>*</sup> |
| c.1429+1G>A<sup>*</sup> | c.2099C>T<sup>*</sup> | c.2206C>T<sup>*</sup> | c.1904G>A<sup>*</sup> | c.1171G>A<sup>*</sup> |
| c.2323-1G>A<sup>*</sup> | c.889T>C<sup>*</sup> | c.2093G>A<sup>*</sup> | c.2102T>A<sup>*</sup> | c.806C>T<sup>*</sup> |

<sup>#</sup> Related to COPD patients<sup>\*</sup> Related to HF patients

**Table S3:** DNA methylation pattern of inflammatory markers in DNMT3A CHIP patients. (**A**) DNA methylation level of inflammatory markers in HF patients with DNMT3A CHIP (n=5) and non-CHIP (n=3). (**B**) DNA methylation level of inflammatory markers in COPD patients with DNMT3A CHIP (n=15) and non-CHIP (n=10) patients.

**A**
| Id | Chr. | Start | End | symbol | entrezID | Methylation level in DNMT3A chip | Methylation level in Non-CHIP | Difference in Methylation level | comb. p.adj.fdr | Combined rank |
| --- | --- | --- | --- | --- | --- | --- | --- | --- | --- | --- |
| ENSG00000136244 | chr7 | 22765503 | 22771621 | *IL6* | 3569 | 0.4453 | 0.446 | -0.001 | 0.487 | 27638 |
| ENSG00000232810 | chr6 | 31543344 | 31546113 | *TNF* | 7124 | 0.462 | 0.4612 | 0.001 | 0.487 | 27078 |
| ENSG00000125538 | chr2 | 113587328 | 113594480 | *IL1B* | 3553 | 0.397 | 0.421 | -0.024 | 0.487 | 4061 |
| ENSG00000169429 | chr4 | 74606223 | 74609433 | *IL8* | 3576 | 0.340 | 0.342 | -0.001 | 0.487 | 28597 |

**B**
| Id | Chr. | Start | End | symbol | entrezID | Methylation level in DNMT3A chip | Methylation level in Non-CHIP | Difference in Methylation level | comb. p.adj.fdr | Combined rank |
| --- | --- | --- | --- | --- | --- | --- | --- | --- | --- | --- |
| ENSG00000136244 | chr7 | 22765503 | 22771621 | *IL6* | 3569 | 0.3958 | 0.4030 | -0.0071 | 0.792 | 22115 |
| ENSG00000232810 | chr6 | 31543344 | 31546113 | *TNF* | 7124 | 0.3986 | 0.4196 | -0.02 | 0.794 | 28298 |
| ENSG00000125538 | chr2 | 113587328 | 113594480 | *IL1B* | 3553 | 0.3454 | 0.3595 | -0.014 | 0.792 | 17032 |
| ENSG00000169429 | chr4 | 74606223 | 74609433 | *IL8* | 3576 | 0.2861 | 0.3015 | -0.015 | 0.669 | 27723 |

**Table S4.** DNA methylation pattern of top differentially methylated genes. The DNA methylation level and P value of top differentially methylated genes between DNMT3A CHIP (HF, n=5; COPD, n=16) and non-CHIP (HF, n=3; COPD, n=10) patients.

| chf_copd_sorted_meth_top_withPvalues |  |  |  |
| --- | --- | --- | --- |
|  | DNMT3A | No_CHIP | pval |
| ENSG00000271119 | 0.0992380640996128 | 0.234585773841018 | 1.89806821253165E-05 |
| SDHAP3 | 0.178225820005913 | 0.299243206206118 | 6.28888826734707E-05 |
| ENSG00000230932 | 0.393408823219886 | 0.468455565050583 | 0.000872656626143761 |
| MUC20-OT1 | 0.312082021439029 | 0.3518119028641 | 0.00159466953986902 |
| SDHAP2 | 0.312082021439029 | 0.3518119028641 | 0.00159466953986902 |
| ENSG00000260774 | 0.144492875081772 | 0.209378814529891 | 0.00177022278637634 |
| SDHA | 0.144492875081772 | 0.209378814529891 | 0.00177022278637634 |
| ENSG00000261235 | 0.887099250130484 | 0.840263177812993 | 0.0021131207135865 |
| GAPDHP49 | 0.136886961065039 | 0.181633452904933 | 0.00362777597874638 |
| ENSG00000263015 | 0.737785874986023 | 0.832959843446773 | 0.0040468301135295 |
| PLEKHH2 | 0.0683426532270443 | 0.099053745118335 | 0.00462208939394681 |
| LCE3C | 0.372957431812154 | 0.521827629353154 | 0.0054941236109396 |
| GSDMD | 0.145531842382765 | 0.166209307128742 | 0.00776008037825875 |
| TUBBP5 | 0.21835106485592 | 0.266357248077789 | 0.00834182469165599 |
| ENSG00000267180 | 0.0802076813672952 | 0.095063896238689 | 0.00837697484722508 |
| ENSG00000270082 | 0.202661941768188 | 0.253441117230957 | 0.00842085666859245 |
| CCDC127 | 0.116757600813096 | 0.163887642932505 | 0.00847219624788187 |
| ENSG00000257281 | 0.129361026036792 | 0.163399008656646 | 0.00952882775398847 |
| LINC01299 | 0.340894808533793 | 0.434176436017962 | 0.0102570881980558 |
| RPL21P111 | 0.876360956366502 | 0.845499371059321 | 0.0106952575848812 |
| ENSG00000229178 | 0.0456570875932378 | 0.0696804826965057 | 0.0109739420122757 |
| ENSG00000263427 | 0.236535494039499 | 0.261516823266601 | 0.0112838627664909 |
| SLC17A9 | 0.259568128071619 | 0.303631528812807 | 0.0115795417262658 |
| LINC01685 | 0.230042504347522 | 0.270723656857061 | 0.0124152274765752 |
| ENSG00000269194 | 0.0829842313178907 | 0.1111406591889 | 0.0126198688132381 |
| LINC02285 | 0.0791245333669025 | 0.0991665239605394 | 0.0131578944941203 |
| ENSG00000237920 | 0.670323469027138 | 0.715012709578997 | 0.0134981428811669 |
| RPSAP17 | 0.670323469027138 | 0.715012709578997 | 0.0134981428811669 |
| ENSG00000257668 | 0.659536880431248 | 0.716241279992801 | 0.0137211319868241 |

